# iDriver: A patient-centric framework for genome-wide cancer driver discovery

**DOI:** 10.64898/2026.02.16.706129

**Authors:** Farideh Bahari, Hesam Montazeri

**Affiliations:** Department of Bioinformatics, Institute of Biochemistry and Biophysics, University of Tehran, Iran

**Keywords:** cancer driver, patient-specific positive selection, mutation heterogeneity, probabilistic graphical model

## Abstract

Tumor genomes harbor a mixture of neutral and positively selected mutations, yet distinguishing true cancer drivers remains a major challenge. Several factors can obscure the detection of selection signals, among which patient-specific variation in mutational burden plays a significant role. Current approaches often fail to account for the heterogeneity in mutation burden across different patients; in particular, no existing method explicitly accounts for it when integrating both mutation recurrence and functional impact. Here we present iDriver, a probabilistic graphical model that integrates both mutation recurrence and functional impact at the individual-patient level, enabling an enhanced estimation of positive selection across functional genomic elements. Applying iDriver to 29 cancer types, we identify both known and previously unrecognized drivers spanning coding and noncoding regions, and provide evidence for their clinical and biological relevance. In comprehensive benchmarks against 12 established driver discovery methods, iDriver consistently outperformed all competitors, achieving the highest rankings for known cancer drivers across both coding and noncoding elements.

## Introduction

Cancer primarily develops through a multistep process of mutation and selection^1^. A small subset of somatic mutations confers selective advantages to affected cells and is referred to as driver mutations^2,3^. Identifying these driver mutations is a central goal in precision oncology, as it enables the development of optimal therapies tailored to a patient’s mutational profile—enhancing treatment efficacy while minimizing toxicity^4–7^. However, distinguishing driver mutations from the more abundant passenger mutations, which do not contribute to tumor progression, remains a major challenge. Driver mutations promote cancer hallmarks such as sustained proliferation, resistance to cell death, and the ability to invade surrounding or distant tissues, and thus are subject to selective pressures^7^. While some recent studies have shown that oncogenes may also exhibit negative selection against loss-of-function mutations^8^, prioritizing candidate cancer drivers often relies on detecting signals of positive selection^7^.

Cancer genomes exhibit substantial heterogeneity across multiple levels—including variation among tissue types, genomic regions, and even among patients with the same cancer type^9,10^. To address this complexity, numerous computational methods have been developed to identify genomic elements that harbor driver mutations by modeling various signals of positive selection while accounting for mutational heterogeneity^7^. These methods can be broadly categorized based on their underlying strategies. Recurrence-based approaches identify regions with a significant excess of mutations by modeling the background mutation rate (BMR)^11–17^. In contrast, functional impact (FI)-based approaches assess whether the observed impact scores of mutations in an element deviate from background expecations—either by comparing them to all possible mutations within the same element or to mutations observed in other elements of the same type^11,12^. Some methods additionally analyze deviations from expected mutational patterns within specific genomic contexts^18^, or leverage metrics such as the ratio of nonsynonymous to synonymous mutations^19,20^. To enhance sensitivity, certain tools combine multiple signals—for example, integrating burden-based models with cancer-type specificity and evolutionary conservation filters^21^. Others incorporate functional consequences without explicit FI scores^21–23^, or merge burden and FI-based analyses^24,25^. To account for heterogeneity across the genome, these methods typically model the BMR using genomic and epigenomic features or local mutation rates, often within machine learning or deep learning frameworks such as Poisson or Negative binomial regression, logistic regression, gradient boosting machines, or convolutional neural networks^23–26^. Tissue-specific heterogeneity is often addressed by fitting cancer-specific models^24,25,27^. However, inter-patient heterogeneity remains largely underexplored—especially in methods combining burden and FI signals to identify coding and non-coding drivers.

Here, we developed iDriver, a probabilistic graphical model that integrates both mutation recurrence and functional impact signals at the patient level, enabling an improved estimation of positive selection across both coding and non-coding genomic elements. To this end, iDriver uses the estimation of background mutation rates obtained from eMET by using 1500 (epi)genomic features affecting mutation rates^28^. It then calculates patient-specific scores of positive selection by quantifying the deviation of observed functional scores from their expected ones considering the patient’s mutational burden. These scores are aggregated and assessed for statistical significance using a null model based on Poisson-Binomial and beta distributions. iDriver can indirectly incorporate other signals of selection pressure, including nucleotide context and clustering of mutations. We applied iDriver to two pan-cancer cohorts and 29 individual cancer types from the PCAWG project. Benchmarking results demonstrated that iDriver outperformed competing methods, achieving the highest rankings for known cancer drivers in both coding and non-coding regions and identifying novel candidate drivers, while maintaining low false positive rates on simulated data. Finally, we highlight and discuss candidate drivers by mapping them to signaling pathways and survival outcomes, and discuss their translational relevance in oncology.

## Results

### Overview of iDriver

We developed *iDriver*, a cancer driver discovery algorithm that identifies driver genomic elements across coding sequences (CDSs), promoters, enhancers, and 5′ and 3′ untranslated regions (UTRs) within specific cohorts (Figure 1). iDriver captures signals of positive selection by integrating mutation recurrence and functional impact, while adjusting for each patient’s overall mutational burden. This adjustment is essential because mutations with a given functional score in high-burden patients are more likely to occur by chance and therefore provide weaker evidence of positive selection than similar mutations in low-burden patients. This signal is often overlooked by recurrence-based driver discovery methods, which generally fail to correct for variability in mutation load across patients.

**Figure 1.**
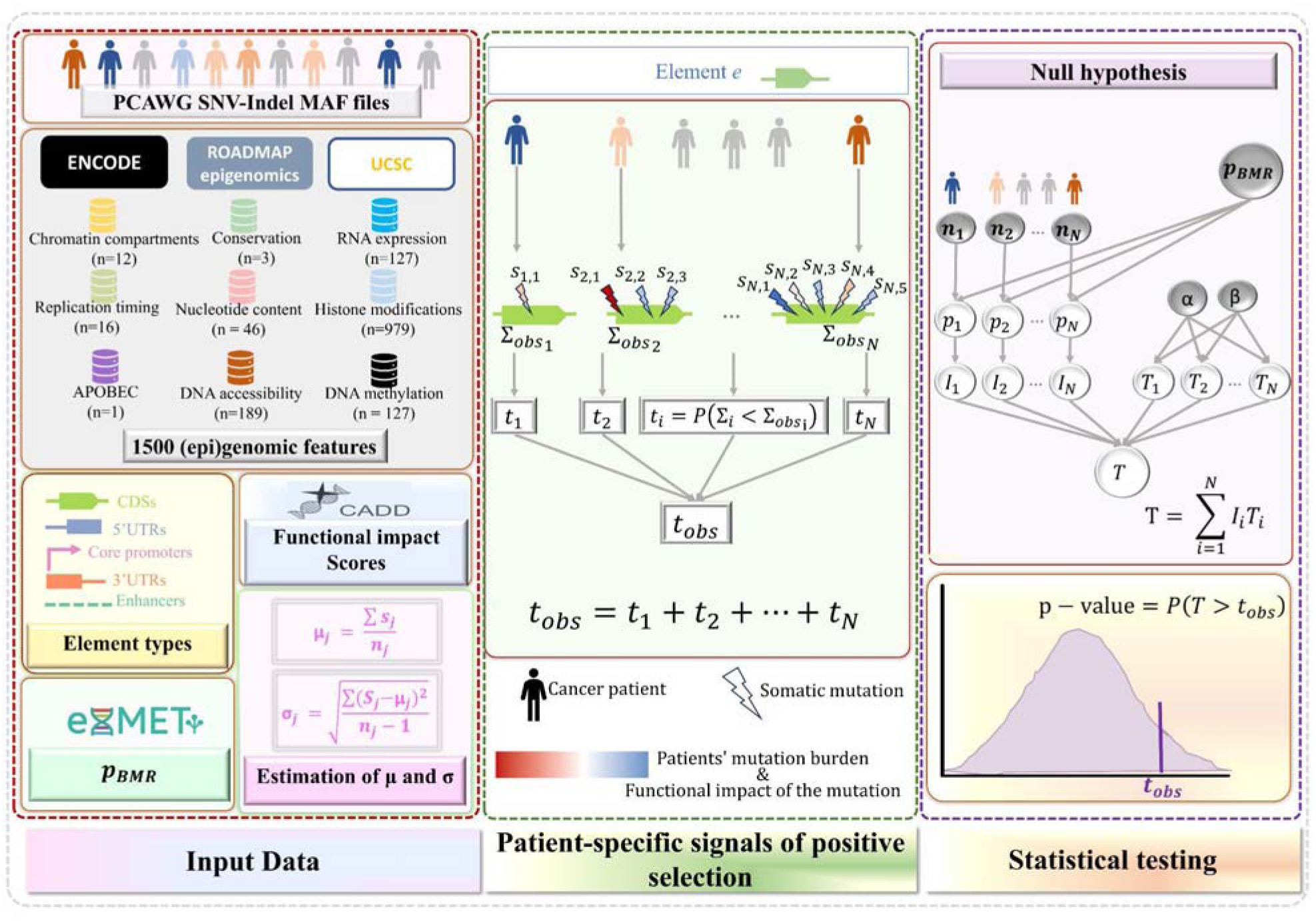
The iDriver algorithm. The iDriver algorithm identifies candidate cancer driver elements by integrating recurrence and functional impact signals at the patient level. The workflow consists of three main steps: 1) BMR estimation: For each genomic element, iDriver estimates a cohort-specific BMR using the eMET model, which incorporates 1,500 (epi)genomic features (e.g., sequence context, replication timing, DNA accessibility, methylation, and APOBEC targets). 2) Calculating patient-specific positive selection signals for each element *e*: For each patient, iDriver calculates a score *t_i_* reflecting the probability that the observed functional impact of a certain number of mutations in a given element exceeds expectation under the null model, adjusting for patient-specific mutation probabilities. Sum of *t_i_*s across all patients in the cohort creates a cohort-level test statistic, *t_obs_* for each element.3) Statistical testing: Observed statistics were compared against a null model of no selection, which accounts for heterogeneous mutation probabilities (Poisson–Binomial), intra-element correlation (linear mixed-effects model), and element length. P-values were then derived by contrasting observed statistic and the null distribution for each element *e*.

To achieve this, iDriver employs a probabilistic graphical model that can be divided into three main steps: (1) Estimating the BMR for each genomic element: Here, we used eMET with 1500 (epi)genomic features—including replication timing, conservation scores, DNA methylation, nucleotide content, epigenomic marks, DNA accessibility, nuclear compartments (HiC), RNA expression, and predicted APOBEC target sites—to estimate the BMR for each genomic element in each cohort^28^. (2) Defining the observed statistic: We quantified patient-specific signals of positive selection of a genomic element by assigning a score *t_i_* to each patient *i*. For a patient with *n* mutations, *t_i_* is defined as the probability that the summation of mutation functional impact scores under the background model exceeds the observed sum. The observed statistic for the given genomic element is then obtained by summing *t_i_* across all patients in the cohort (see Methods for details). iDriver can incorporate any functional impact scoring system, in this study we used CADD scores^29^. (3) Statistical testing: we calculated a p-value for each genomic element by comparing the observed statistic to the null model, i.e. *P*(*T* > *t_obs_*). The null model, represented by a probabilistic graphical model (Figure 1; see Methods for details), characterizes the distribution of the test statistic *T* under background model. The conditional tail probability *P*(*T* > *t_obs_*) was used to assess whether a given element is a driver. This probability was approximated using a normal distribution, with the mean and variance calculated from the Beta distribution of patient-level scores. To account for intra-class correlation among mutations within the same element, we applied a linear mixed-effects model, stratified by element length. In addition, a Poisson–Binomial distribution was used to model heterogeneous mutation probabilities across patients. Therefore, in iDriver, we aimed to capture three levels of heterogeneity: (i) across cohorts, by estimating distinct BMR models for each cancer cohort; (ii) across genomic regions, by incorporating a comprehensive set of features that influence mutation rates and (iii) across patients, by refining the probability of observing mutations in each patient according to their total mutational burden.

### iDriver outperforms other driver discovery methods

We applied iDriver to two pan-cancer cohorts of non-hypermutated donors, as well as to 29 cancer-specific cohorts. The first pan-cancer cohort, referred to as the Pancancer cohort, includes 2,514 non-hypermutated donors across 37 cancer types. The second, the Pancancer* cohort, comprises 2,253 donors from the Pancancer cohort, excluding those with skin melanoma and lymphoma.

Mutation burden differs markedly across cohorts and among donors, highlighting inter-patient heterogeneity (Figure 2a). Among all donors, 69 were hypermutated, exhibiting more than 30 mutations per megabase—primarily from the Skin-Melanoma and Colorectal-AdenoCA cohorts. Such hypermutated genomes were excluded in previous studies to facilitate more accurate BMR estimation^23,24^. Beyond inter-patient variability, distinct genomic elements also display heterogeneous mutation rates and functional impacts. The top 60 most mutated coding and noncoding elements, showing observed versus expected mutation counts estimated by eMET, along with their distinct functional effects for the Pancancer* cohort are provided in Figure 2b, and Supplementary Figure 1, respectively. Notably, elements exhibiting high mutational burden and/or strong high functional impact correspond to well-established cancer genes, which are involved in main cancer pathways (Figures 2c,d).

**Figure 2.**
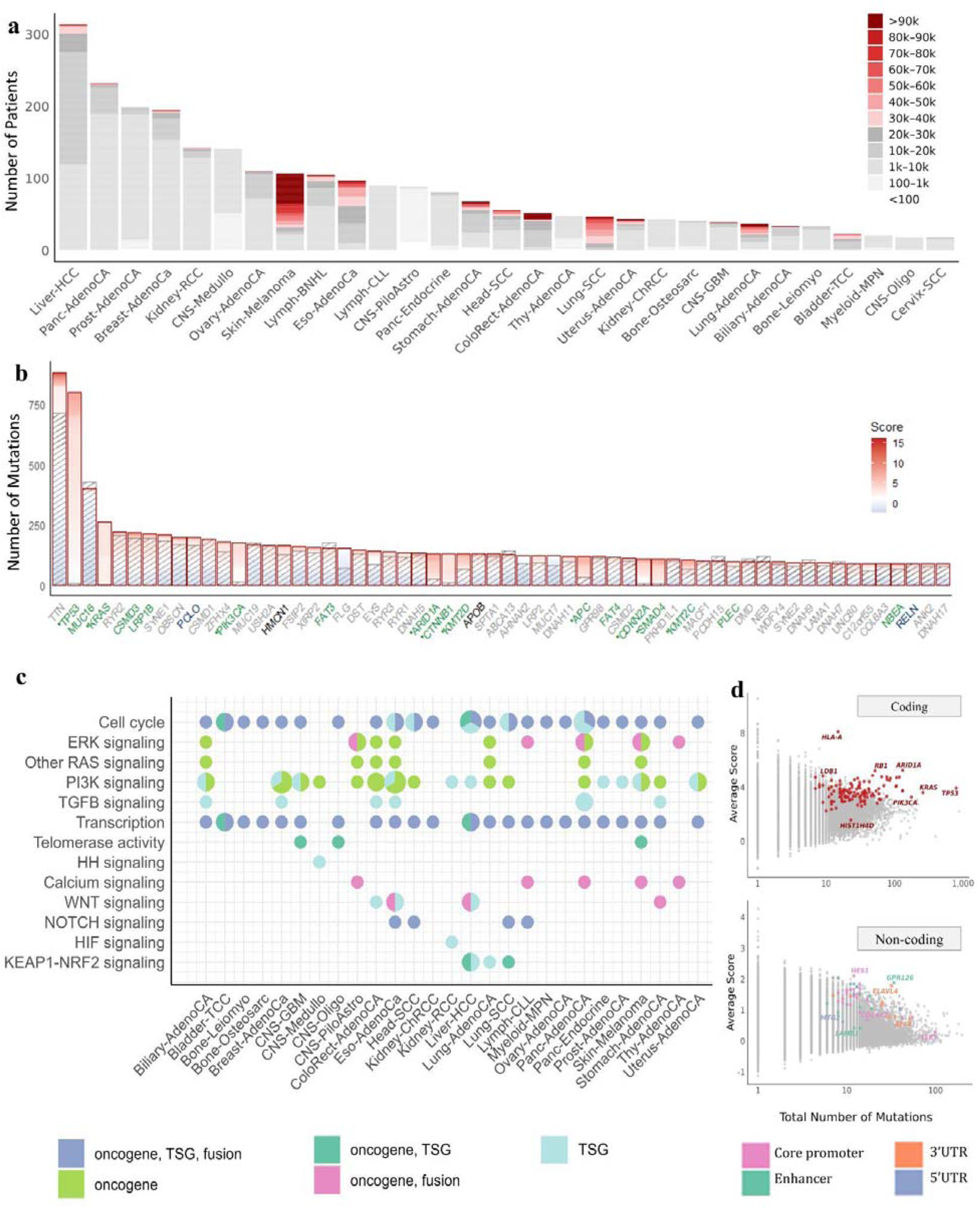
Summary of the data and distribution of known cancer genes reidentified by iDriver across cancer-pathways. **a** Heterogeneity across cohorts and patients is depicted as a stacked bar plot. Each bar represents a cohort, with patients stacked according to their mutation burdens. Within each stack, darker reds indicate higher tumor burden. Skin-Melanoma and Colorectal-AdenoCA show the largest numbers of hypermutated cases. **b** The top 60 most frequently mutated CDSs, showing both observed (highlighted by red bars) and expected numbers of mutations estimated using eMET (indicated by diagonal stripes in the bars). Mutations within each element are represented as stacked bars, colored according to their CADD pathogenicity scores. Gene symbol colors indicate whether the gene is listed as a CGC gene (green), an OncoKB cancer gene (blue), a PCAWG cancer gene (black), or not listed in any cancer gene database (grey). Significant hits identified by iDriver are highlighted with an asterisk beside the gene name. **c** Mutated pathways (KEGG cancer networks) across cancer cohorts are shown as scatter pie charts. For each cohort–pathway pair, the size of the pie chart represents the number of mutated genes, and the segments indicate the distribution of these genes according to their annotated roles in cancer. This plot shows that several pathways are recurrently mutated in multiple cancer types. **d** Total number of mutations in each coding and non-coding element versus the average CADD score of mutations within that element is displayed as scatter plots. Highlighted points indicate some of the significant hits identified by iDriver. For non-coding elements, significant hits are colored according to element type (3′UTR, 5′UTR, enhancer, or core promoter).

Among the top 60 most frequently mutated coding genes are *TP53*, *KRAS*, *PIK3CA*, *ARID1A*, and *CTNNB1*, all are recurrently mutated in cancer. For some genes like *HLA-A* we observed 15 mutations among 2253 patients—although exceeding its expected rate—it ranked among the highest in terms of average functional impact scores (Figure 2c).

We benchmarked iDriver against 12 published methods across all 31 cohorts, directly adopting the published p-values reported by the PCAWG Drivers and Functional Interpretation Working Group (PDFIWG)^9^. These methods—each capturing signals of positive selection from a distinct perspective—were developed to identify both coding and non-coding driver elements. Some methods such as NBR^14^, ExInAtor^15^, ncdDetect^16^ and LARVA^13^ are burden-based approaches. oncodriveFML^11^ (with CADD and VEST scores) relies solely on functional impact testing, while ncDriver^21^ combines a burden-based model with cancer specificity and conservation filters. MutSig^22^ and ActiveDriverWGS^23^ are primarily burden-based but incorporate functional impacts without explicitly using the corresponding scores. Other approaches include regDriver^30^, dNdScv^19^, and CompositeDriver^12^. CompositeDriver was excluded from our benchmark due to its reliance on FunSeq scores^31^, which may introduce information leakage when evaluating mutations in known cancer genes (Supplementary Note 1). Finally, DriverPower combines both burden- and FI-based signals at the population level.

We defined the genes in the Cancer Gene Census as the gold standard, and any significant hits within this set were considered true positives (TPs). Benchmarking showed that iDriver was the top-performing method for non-coding driver discovery in 22 of 27 cohorts (81.5%), compared with MutSig and ncdDetect, which each led in 2 cohorts (7% each). For coding element prioritization, iDriver ranked first in 12 of 27 cohorts (44.4%), followed by DriverPower in 7 cohorts (26.9%) and dNdScv in 4 cohorts (15.4%) (Figure 3 and Supplementary Figs. 2-3). This further highlights the value of combined approaches and, in particular, the importance of accounting for inter-patient heterogeneity in mutation rates—an area where iDriver demonstrates a clear advantage over DriverPower and MutSig. Unless otherwise specified, we applied iDriver to non-hypermutated donors, pooling all mutation types including SNVs, DNVs, MNVs, and indels to estimate the BMR. Nonetheless, we also demonstrated its superiority in cohorts containing both non-hypermutated and hypermutated patients (Supplementary Figures 4–6). In addition, we performed dedicated analyses on a cohort composed exclusively of hypermutated donors, explored the signals of positive selection using iDriver based solely on synonymous mutations, and separately modeled SNVs and non-SNVs, subsequently integrating their p-values using different combination strategies. The results of these analyses are provided in Supplementary Note 2–4, Supplementary Figures 7–9, and Supplementary Data 1, 2.

**Figure 3.**
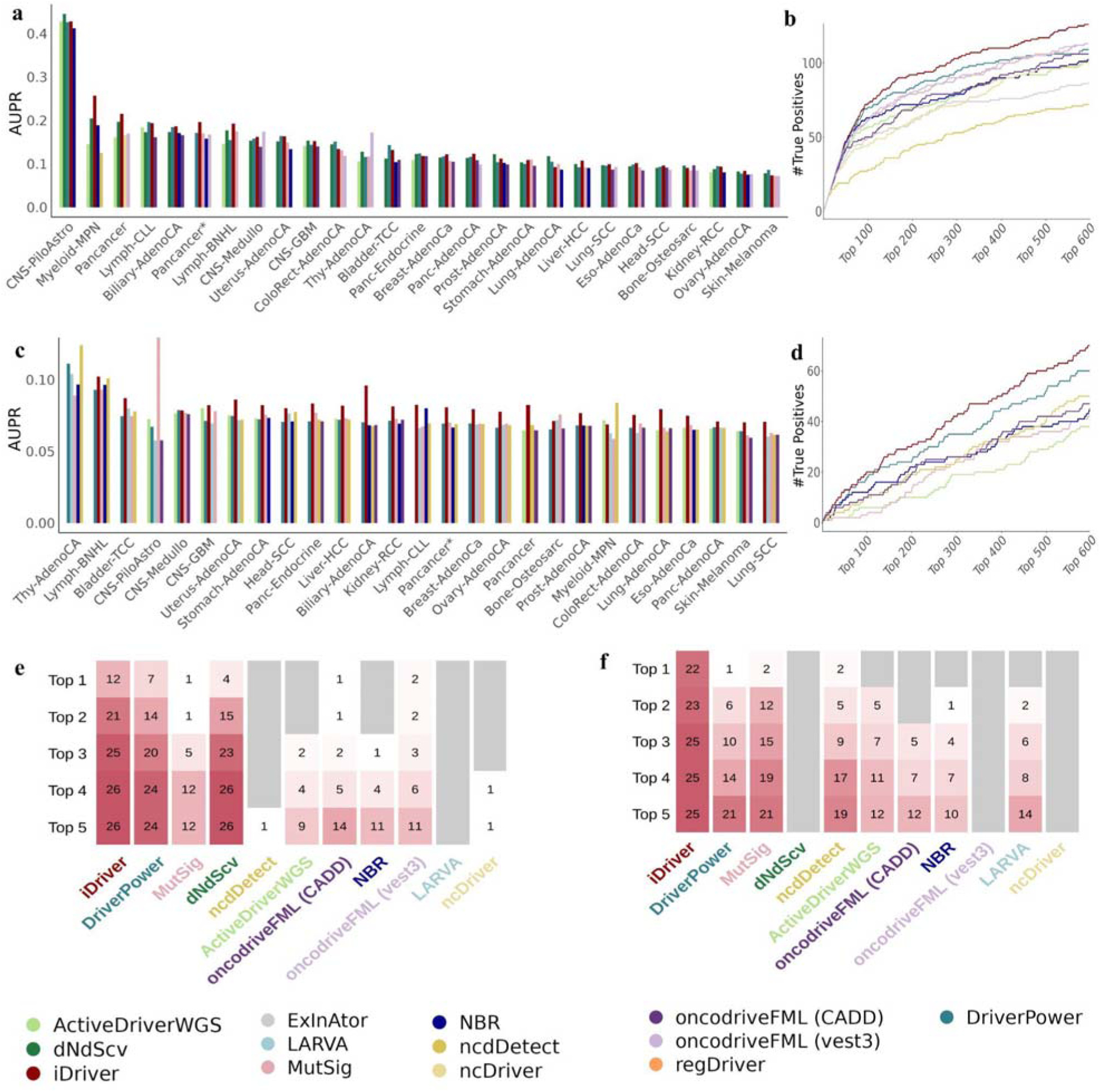
Benchmarking iDriver against 12 driver discovery methods. Performance comparison of iDriver with 12 published driver discovery methods across 31 PCAWG cancer cohorts for both coding (a, b, and e panels) and non-coding (c, d and f panels) elements. a and c panels: AUPR per cancer cohort for top 5 methods. b and d panels: Cumulative number of TPs among the top-ranked elements (up to top 600) for each method for the Pancancer* cohort, demonstrating the superior ranking ability of iDriver. The heatmaps in panels e and f summarize the results shown in panels a and c, showing how often each method ranks among the top-k methods for driver discovery based on AUPR across all cohorts. Overall, this figure shows iDriver consistently outperformed other methods, including both recurrence-based, functional-impact-based, and combined approaches, by preserving the best ranking of the cancer drivers. True positives were defined as elements overlapping with Cancer Gene Census genes.

### Pan-Cancer Identification of Coding and Non-Coding Drivers

In the pan-cancer cohort, iDriver identified 346 significant elements at a 10% false discovery rate (FDR < 0.1), including 214 coding and 132 non-coding candidates. More than two-thirds of these correspond to established cancer genes cataloged in COSMIC, OncoKB, or PCAWG, while the remainder represent previously unknown candidates with potential biological relevance. In the more stringent pancancer* cohort, which excludes Skin-melanoma, Lymphomas, and hypermutated donors, 176 elements were significant (125 coding and 51 non-coding), among which 39 (22 coding and 17 non-coding) were not previously annotated in known cancer gene databases (Supplementary Data 3).

Among coding elements, all of the top 63 significant hits in the pancancer* cohort correspond to known drivers, including *TP53*, *KRAS*, *PIK3CA*, *CDKN2A*, *VHL*, and *CTNNB1*. *SMARCA4* ranked 63rd among coding regions, consistent with its known role as a tumor suppressor in non–small-cell lung and ovarian cancers^32^. The first candidate coding driver not previously reported in cancer gene databases was *LRP12* (64th among coding and 90th overall), followed by *REM1*, *PAF1*, *HNRNPK*, *PROS1*, *MEIS2*, *RPL11*, and *ATP1A2*—all within the top 100 coding hits.

Pathway-level analysis revealed a convergence of iDriver-identified drivers on signaling and regulatory circuits associated with tumor progression (Figure 4a–b). In particular, the PI3K–AKT, MAPK signaling pathways and cellular senescence, along with infection-related pathways such as human papillomavirus and Kaposi sarcoma–associated herpesvirus infection, harbored the largest numbers of significant drivers (Supplementary Figures 10,11). Several candidate genes were enriched across multiple cancer-relevant pathways (e.g., *PIK3CA*, *HRAS*, *KRAS*, and *NRAS* each appeared in more than 15 KEGG pathways). Figure 4b highlights representative pathways for each gene, while Supplementary Figure 11 provides a more comprehensive list of mutated pathways for each gene. Notably, several infection-related KEGG pathways include key cancer-associated processes—such as Cell cycle, PI3K signaling, Apoptosis, and Wnt signaling—making them among the most frequently mutated pathways across multiple driver candidates.

**Figure 4.**
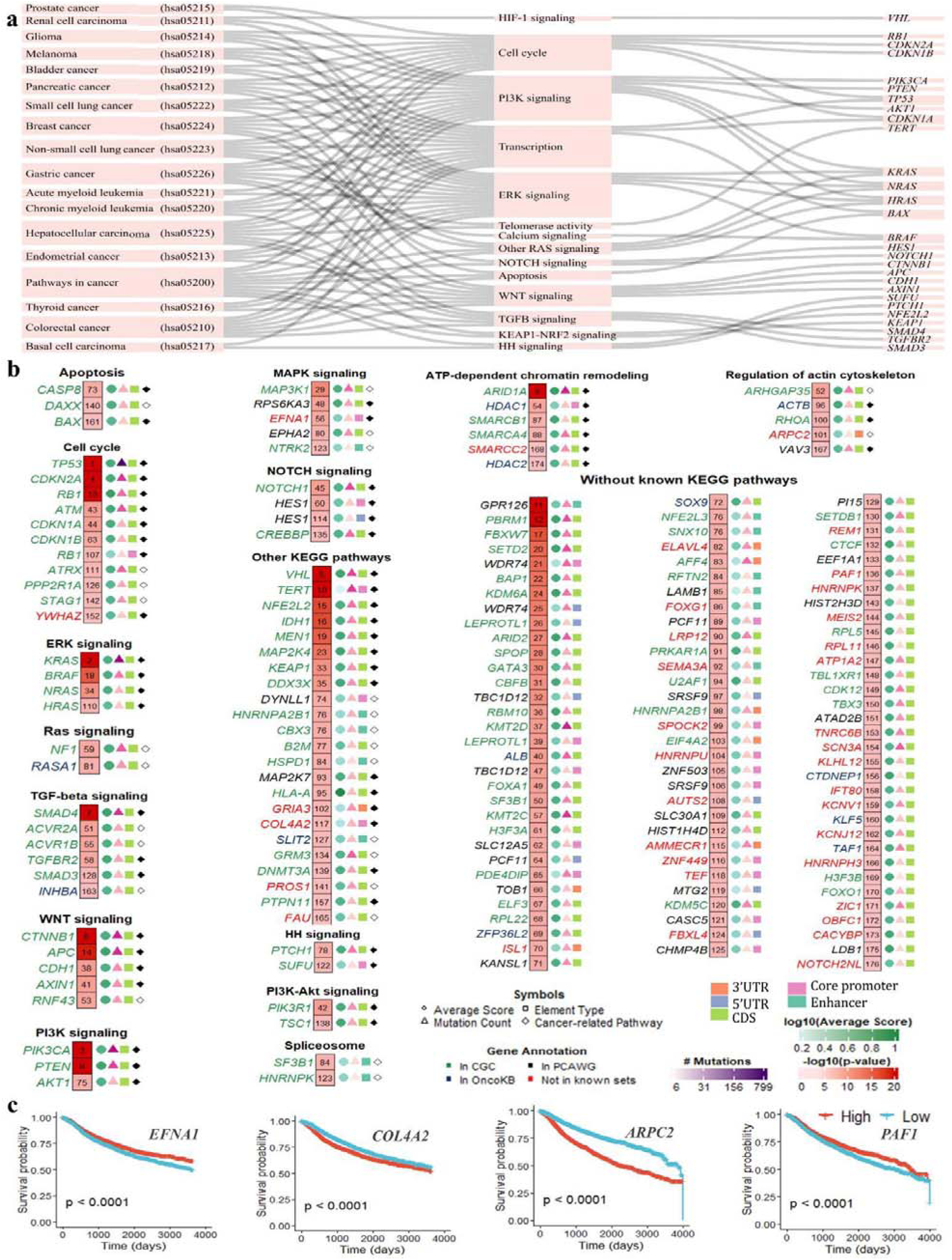
Pancancer* candidate drivers identified by iDriver. **a** Mutated KEGG cancer pathways and their networks associated with iDriver-significant hits. **b** This plot shows all candidate drivers detected by iDriver, ranked by statistical significance and grouped according to their associated biological pathway. The value inside each cell indicates the candidate’s rank among all coding and non-coding hits for the Pancancer* cohort. The three colored dots to the right of each cell represent, from left to right: the average CADD score of mutations in the element, the total number of mutations in the element, and the element type. Gene name colors indicate prior knowledge: Green – listed in the COSMIC CGC, Blue – listed in OncoKB, Black – listed in the PCAWG gene list. Red gene names indicate candidates not in any of the above resources. Black-filled diamonds indicate cancer genes identified in the same cancer type according to the CGC. Empty diamonds indicate genes classified as cancer drivers in other cancer types. **c** Kaplan–Meier plot comparing survival probability between samples with high (red) versus low (blue) expression, using data from UCSC Xena.

Finally, several of the newly identified drivers, including *EFNA1*, *COL4A2*, and *ARPC2*, were associated with reduced patient survival across multiple cohorts (Figure 4c), supporting their potential clinical significance. Literature evidence for novel candidate drivers is summarized in Table 1. Results from applying iDriver to two additional pan-cancer cohorts—including (i) all donors (both hypermutated and non-hypermutated) and (ii) only hypermutated donors—are available in the Supplementary Data 4, 5.

**Table 1.**
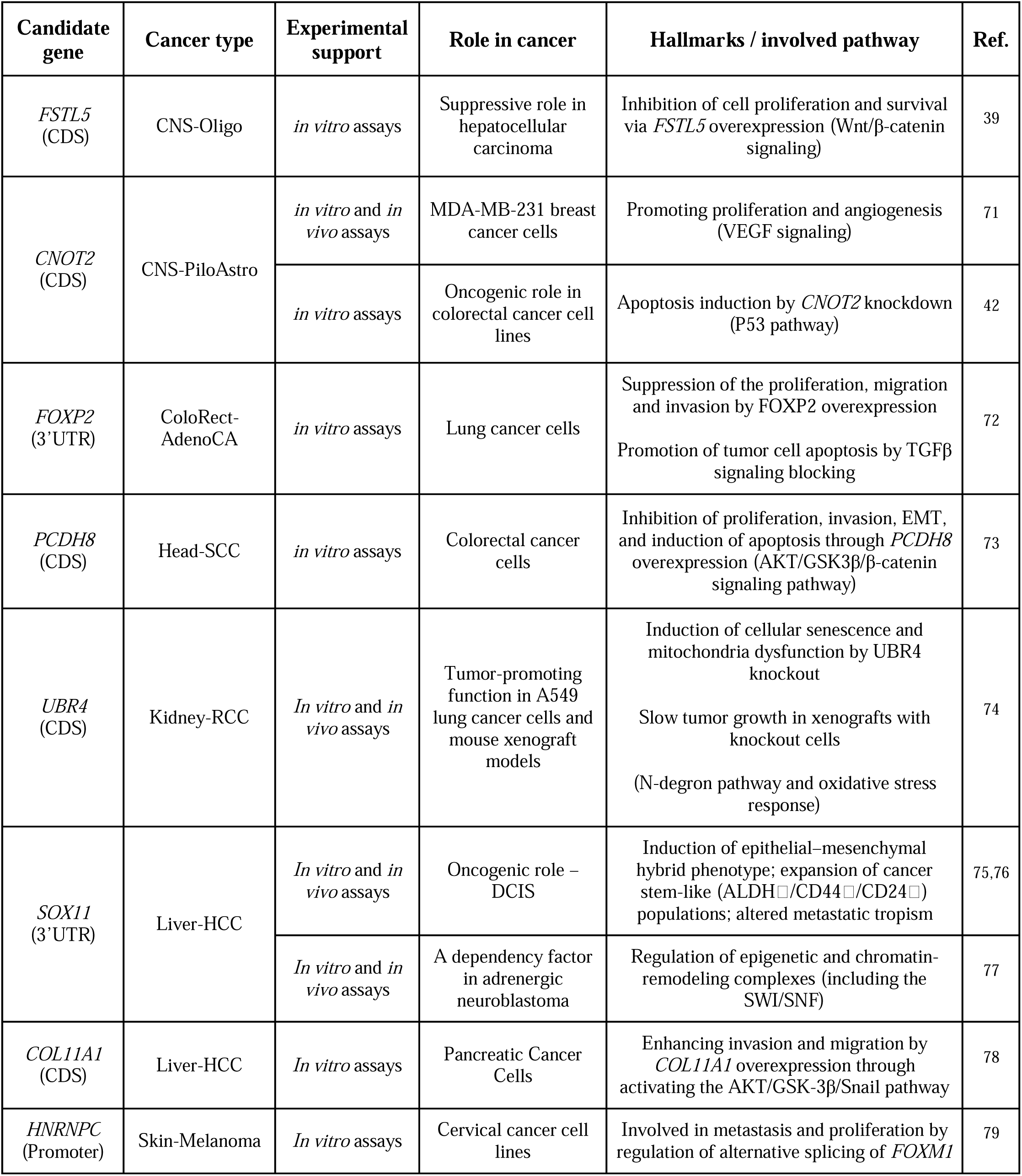
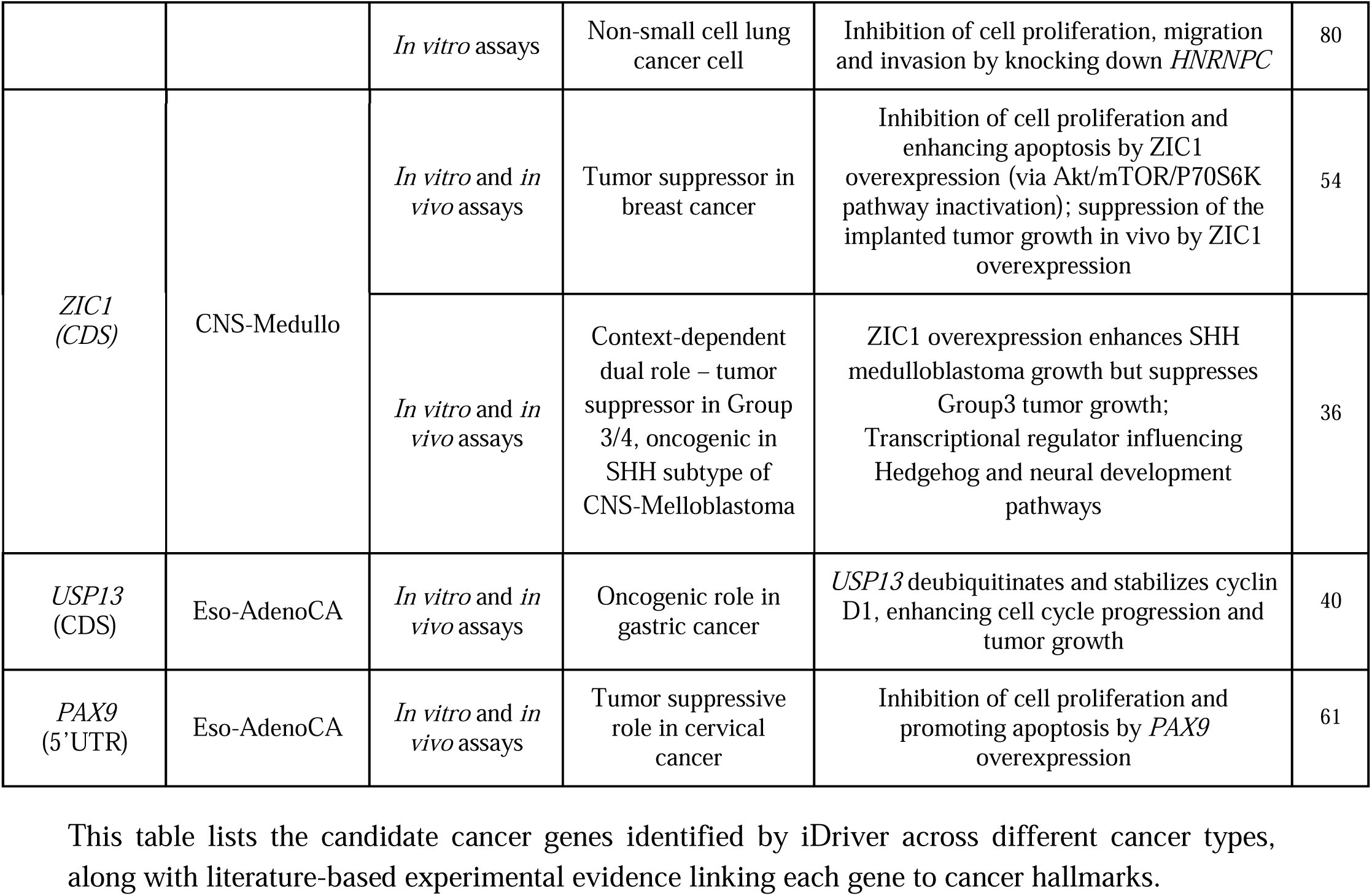
Selected candidate cancer genes identified by iDriver.

| Candidate gene | Cancer type | Experimental support | Role in cancer | Hallmarks / involved pathway | Ref. |
| --- | --- | --- | --- | --- | --- |
| *FSTL5* (CDS) | CNS-Oligo | *in vitro* assays | Suppressive role in hepatocellular carcinoma | Inhibition of cell proliferation and survival via *FSTL5* overexpression (Wnt/β-catenin signaling) | 39 |
| *CNOT2* (CDS) | CNS-PiloAstro | *in vitro* and *in vivo* assays | MDA-MB-231 breast cancer cells | Promoting proliferation and angiogenesis (VEGF signaling) | 71 |
|  |  | *in vitro* assays | Oncogenic role in colorectal cancer cell lines | Apoptosis induction by *CNOT2* knockdown (P53 pathway) | 42 |
| *FOXP2* (3'UTR) | ColoRect-AdenoCA | *in vitro* assays | Lung cancer cells | Suppression of the proliferation, migration and invasion by *FOXP2* overexpression<br>Promotion of tumor cell apoptosis by TGFβ signaling blocking | 72 |
| *PCDH8* (CDS) | Head-SCC | *in vitro* assays | Colorectal cancer cells | Inhibition of proliferation, invasion, EMT, and induction of apoptosis through *PCDH8* overexpression (AKT/GSK3β/β-catenin signaling pathway) | 73 |
| *UBR4* (CDS) | Kidney-RCC | *In vitro* and *in vivo* assays | Tumor-promoting function in A549 lung cancer cells and mouse xenograft models | Induction of cellular senescence and mitochondria dysfunction by *UBR4* knockout<br>Slow tumor growth in xenografts with knockout cells<br>(N-degron pathway and oxidative stress response) | 74 |
| *SOX11* (3'UTR) | Liver-HCC | *In vitro* and *in vivo* assays | Oncogenic role – DCIS | Induction of epithelial–mesenchymal hybrid phenotype; expansion of cancer stem-like (ALDH$^+$/CD44$^+$/CD24$^-$) populations; altered metastatic tropism | 75,76 |
|  |  | *In vitro* and *in vivo* assays | A dependency factor in adrenergic neuroblastoma | Regulation of epigenetic and chromatin-remodeling complexes (including the SWI/SNF) | 77 |
| *COL11A1* (CDS) | Liver-HCC | *In vitro* assays | Pancreatic Cancer Cells | Enhancing invasion and migration by *COL11A1* overexpression through activating the AKT/GSK-3β/Snail pathway | 78 |
| *HNRNPC* (Promoter) | Skin-Melanoma | *In vitro* assays | Cervical cancer cell lines | Involved in metastasis and proliferation by regulation of alternative splicing of *FOXM1* | 79 |
|  |  | *In vitro* assays | Non-small cell lung cancer cell | Inhibition of cell proliferation, migration and invasion by knocking down *HNRNPC* | 80 |
| *ZIC1*<br>(CDS) | CNS-Medullo | *In vitro* and *in vivo* assays | Tumor suppressor in breast cancer | Inhibition of cell proliferation and enhancing apoptosis by *ZIC1* overexpression (via Akt/mTOR/P70S6K pathway inactivation); suppression of the implanted tumor growth *in vivo* by *ZIC1* overexpression | 54 |
|  |  | *In vitro* and *in vivo* assays | Context-dependent dual role – tumor suppressor in Group 3/4, oncogenic in SHH subtype of CNS-Medulloblastoma | *ZIC1* overexpression enhances SHH medulloblastoma growth but suppresses Group3 tumor growth; Transcriptional regulator influencing Hedgehog and neural development pathways | 36 |
| *USP13*<br>(CDS) | Eso-AdenoCA | *In vitro* and *in vivo* assays | Oncogenic role in gastric cancer | *USP13* deubiquitinates and stabilizes cyclin D1, enhancing cell cycle progression and tumor growth | 40 |
| *PAX9*<br>(5'UTR) | Eso-AdenoCA | *In vitro* and *in vivo* assays | Tumor suppressive role in cervical cancer | Inhibition of cell proliferation and promoting apoptosis by *PAX9* overexpression | 61 |

This table lists the candidate cancer genes identified by iDriver across different cancer types, along with literature-based experimental evidence linking each gene to cancer hallmarks.

### Tissue-Specific Discovery of Cancer Drivers

Applying iDriver to 29 cancer-specific cohorts identified 415 significant coding and non-coding elements (q < 0.1), of which 362 overlapped with known cancer genes reported in COSMIC CGC, OncoKB, or PCAWG (Figure 5a; Supplementary Data 3). These results confirm iDriver’s ability to recover well-established drivers while also revealing previously unrecognized candidates with potential tissue-specific relevance.

**Figure 5.**
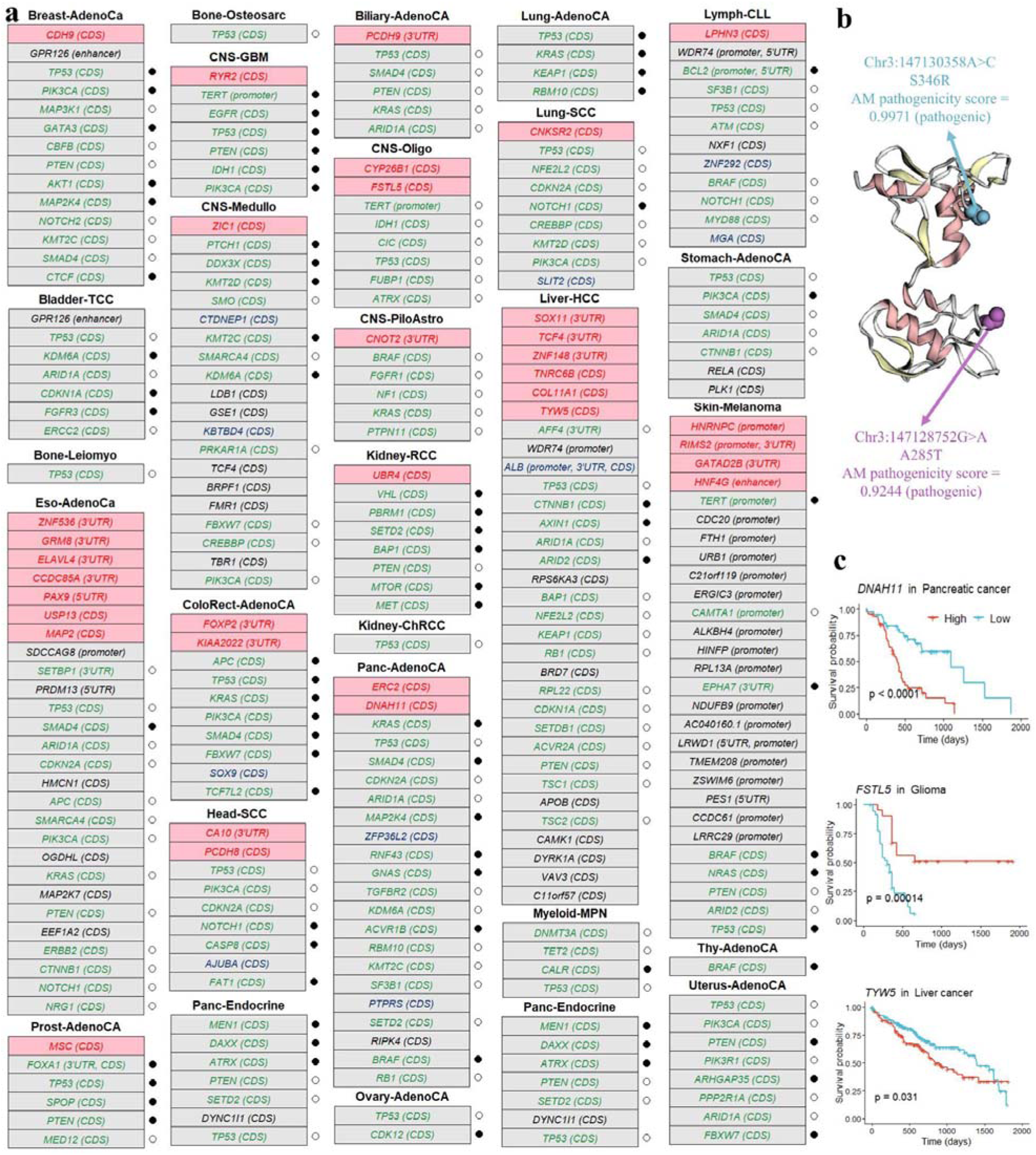
Coding and non-coding candidate drivers identified by iDriver across cancer cohorts. **a** Driver candidates detected by iDriver, grouped by cancer type. Genes highlighted at the top of each cancer type represent novel hits identified by iDriver. Gene name colors indicate prior evidence: Green: reported in the COSMIC CGC, Blue: listed in OncoKB, Black: included in the PCAWG cancer gene list, and Red: candidates not reported in previous cancer gene lists. For CGC genes, the circles to the left of each table indicate whether the gene has been previously reported for the same cancer type in COSMIC CGC (black-filled circles) or for a different cancer type (empty circles). **b** Example visualization of SNV mutations in the *ZIC1* CDS using the Mutation Position Imaging Toolbox (MuPIT), annotated with alpha missense pathogenicity scores. **c** Kaplan–Meier plot comparing survival probability between samples with high (pink) versus low (blue) expression, using data from the Human Protein Atlas.

Across cohorts, top-ranking elements corresponded to canonical cancer genes such as *TP53*, *KRAS*, *PIK3CA*, and *ARID1A*, reflecting major oncogenic processes including Cell cycle, apoptosis, cellular senescence and MAPK and PI3K signaling (Supplementary Figure 12). Some pathways, such as Cell Cycle and PI3K signaling, were altered across most cancer types, while others displayed clear tissue specificity. Notably, HIF signaling was predominantly affected in Kidney-RCC, consistent with frequent *VHL* inactivation leading to constitutive stabilization of HIF-1α and HIF-2α, a hallmark of renal tumorigenesis^33^. In contrast, cGMP–PKG signaling was mutated almost exclusively in Breast-AdenoCA. Although canonical cGMP–PKG genes were not directly mutated, alterations in *AKT1*—a component of the PI3K–AKT pathway—and its downstream shared targets contribute to cGMP–PKG pathway perturbation in this cohort (Figure 2c, Supplementary Figure 12).

Beyond recovering known drivers, iDriver uncovered several coding-region candidates not previously linked to cancer, such as *ZIC1* in CNS-Medullo, *FSTL5* in CNS-Oligo, *CDH9* in Breast-AdenoCA, *RYR2* in CNS-GBM, *TYW5* and *COL11A1* in Liver-HCC, and *DNAH11* in Pancreatic-AdenoCA. These genes may represent context-dependent drivers that operate within distinct tissue environments. Among non-coding elements, iDriver identified several regulatory regions with potential functional impact, including the *PAX9* 5′UTR in Eso-AdenoCA, and the *HNRNPC* promoter and *HNF4G* enhancer in Skin-Melanoma. In total, 14 3′UTR candidates were uniquely discovered, such as *CNOT2* in CNS-PiloAstro and *FOXP2* in ColoRect-AdenoCA.

Survival analyses using the Human Protein Atlas revealed strong prognostic associations for *DNAH11* in pancreatic cancer (Figure 5c). The rs2285947 polymorphism in *DNAH11* has been linked to an elevated risk of cancer^3435^. Other examples of genes with strong prognostic value include *TYW5* in Liver-HCC and *FSTL5* in glioma (Figure 5c). Notably, *FSTL5* has strong literature support connecting it to several cancer hallmarks (see Table 1 and the *Functional and Literature Support for Candidate Drivers* section for details). Interestingly, *ZIC1* was identified by iDriver as a cancer driver in CNS-Medullo, and recent functional studies support its role as a context-dependent driver in medulloblastoma^36^. Structural mapping of *ZIC1* coding variants using MuPIT revealed mutations enriched for high AlphaMissense pathogenicity scores, further supporting their potential functional relevance (Figure 5b).

### False Positive Rate Evaluation Using Simulated Mutation data

We evaluated the iDriver algorithm for its type I error rate. To this end, we applied iDriver to non-hypermutated samples from the Sanger-simulated dataset generated by the PDFIWG, covering 29 cancer types and two pan-cancer cohorts, and adjusted the p-values for controlling false discovery rate, and reported number of significant elements across each cancer type (q-value < 0.1). The analysis revealed no false positives (FPs) in the cancer-specific assessments, except for the Lymph-BNHL cohort, which showed 11 FPs—likely due to AID off-target effects and the resulting localized hypermutation. The pan-cancer cohort reported 9 FPs, whereas the pancancer* cohort showed no FPs (Supplementary Data 6). Comparison of FP rates across cancer types and PCAWG methods revealed that for Lymph-BNHL, LARVA, NBR, and MutSig reported more false positives than iDriver, whereas ActiveDriverWGS did not report any FPs (Supplementary Figure 13). The number of false positives introduced by iDriver and previous driver discovery methods after multiple-testing correction, separately for coding and non-coding elements across all cohorts, are shown in Figure 6a.

**Figure 6.**
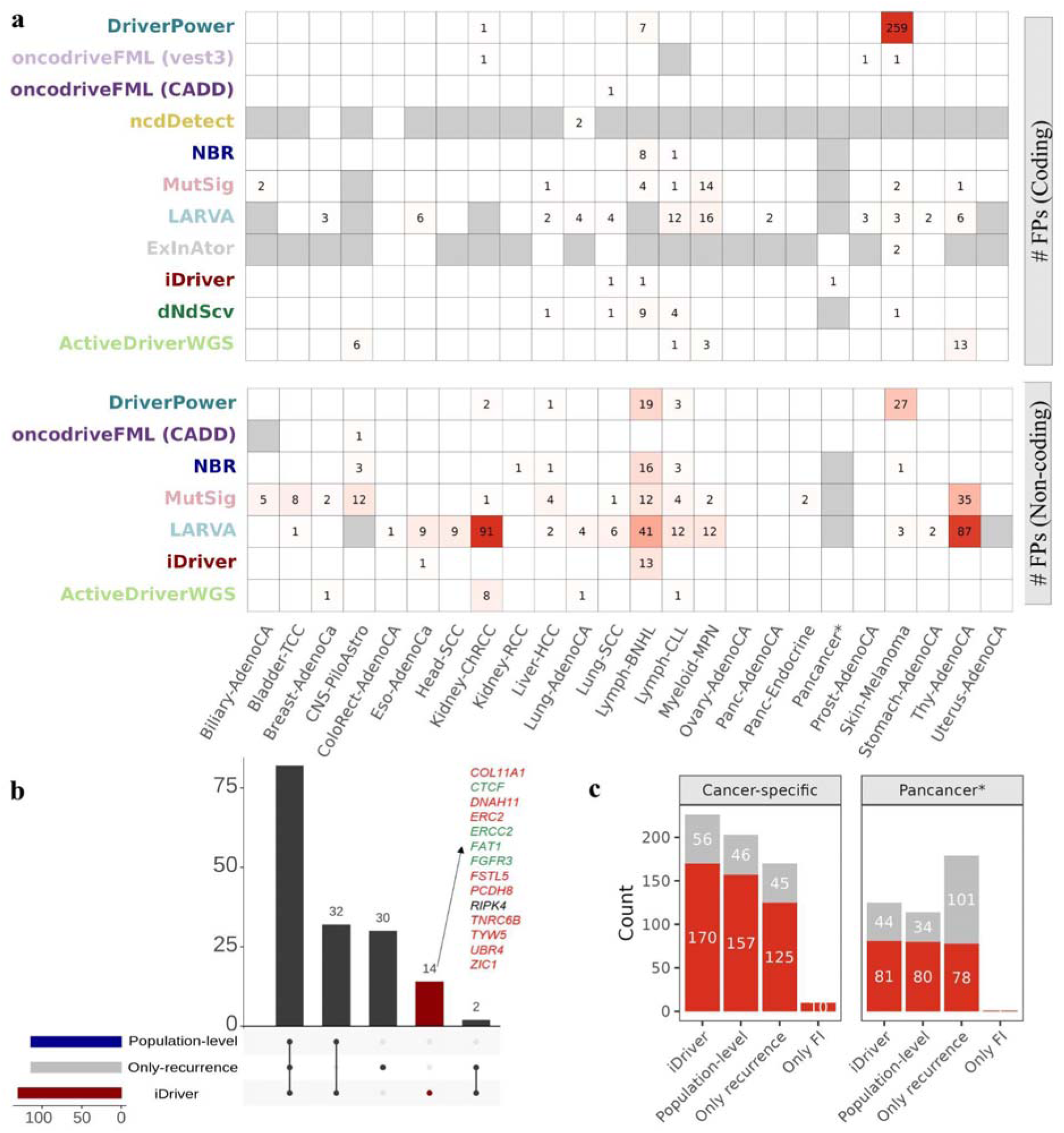
iDriver ablation study results, and False discovery rate analysis. Panel a shows a heatmap of the number of false positives reported for the simulated data across cohorts by different methods. White cells indicate that the method did not report any false positives for a given cohort. Grey cells indicate that either no results were available for the method in that cohort, or that the cohort had reported p-values for fewer than 95% of the elements compared to the union of iDriver and DriverPower, making the comparison unfair. Panel b shows the number of coding cancer genes identified by iDriver for cancer-specific cohorts and each of the ablated iDriver models, highlighting the number of shared and uniquely identified genes by each method. Genes exclusively identified by iDriver are shown separately. Gene name colors indicate prior evidence: Green = reported in COSMIC CGC, Black = included in the PCAWG cancer gene list, and Red = candidates not reported previously. Panel c shows the number of true positives and the number of significant hits not reported in the COSMIC CGC for the original iDriver (iDriver) and three ablated component settings for coding elements. For the cancer-specific block of panel c, the plot presents the sum of true positives and non-CGC candidates across different cancer types, excluding Lymph-BNHL.

### Influence of ablating the components of iDriver on performance

Given the improved performance of iDriver in the benchmark across different cohorts, we performed an ablation study. To this end, we compared the performance of iDriver in its original configuration— which accounts for heterogeneity in mutation rates across individual patients by incorporating recurrence, functional impact scores, and intracorrleation of scores within each element—with four ablated component settings: (i) the population-level setting, (ii) the uncorrelated score setting, (iii) recurrence only, and (iv) functional impact only (FI only). In the population-level setting, each patient was assigned the same total number of mutations, equal to the average total mutations observed in the cohort, and iDriver was run with its default parameters. The original, individual-level configuration yielded a larger number of significant hits without a substantial increase in type I error compared to the population-level model. All hits identified in the population-level analysis were also captured by the individual-level approach; however, the latter reported an additional 103 significant elements across all cohorts. Among these, 41 were annotated as cancer genes in at least one of the CGC COSMIC, OncoKB, or PCAWG cancer gene lists. Examples include CDSs of *FGFR3* and *ERCC2* in Bladder-TCC, *CTCF* in Breast-AdenoCa, *PIK3CA* in CNS-GBM, *CTNNB1*, *NOTCH1*, and *NRG1* in Eso-AdenoCa, *FAT1* in Head-SCC, and *RIPK4*, *BRAF*, and *RB1* in Panc-AdenoCa. Notably, several novel candidates were exclusively detected in the individual-level setting. These include: Coding sequences of *ZIC1* in CNS-Medullo, *FSTL5* in CNS-Oligo, *MAP2* in Eso-AdenoCa, *PCDH8* in Head-SCC, *UBR4* in Kidney-RCC, *TNRC6B*, *COL11A1*, and *TYW5* in Liver-HCC, and *ERC2* and *DNAH11* in Panc-AdenoCa. And 3′UTRs of *PCDH9* in Biliary-AdenoCa, *GATAD2B* and *RIMS2* in Skin-Melanoma, *ZNF148* and *TCF4* in Liver-HCC, *CA10* in Head-SCC, and the enhancer of *HNF4G* in Skin-Melanoma. These findings demonstrate that incorporating patient-specific mutational heterogeneity into iDriver’s framework substantially improves its sensitivity for detecting both known and novel cancer drivers while maintaining control over false positive rates.

Since certain genomic regions are reported to have higher mutation rates due to their genomic and epigenomic features^24,26,37^ and/or to harbor variants with higher CADD scores^27^, we accounted for the intracorrelation of observed statistics within each element under study. This was done across elements of the same type, grouped into the same length category. This approach not only considers the correlation between mutation counts and the accumulation of variants with similar CADD scores in a genomic region, but also indirectly accounts for deviations from the null model that could indicate hotspot formation within an element. To evaluate the impact of incorporating intracorrelation, we compared iDriver’s performance with and without considering intracorrelation of the observed statistics. When iDriver was run on Sanger simulated data, under the assumption that the observed statistics of an element are uncorrelated (Rho = 0), it produced a large number of false positives—1,421 and 1,095 significant hits in the simulated data for the Pan-cancer and Pancancer* cohorts, respectively. In contrast, incorporating intracorrelation reduced false positives dramatically, to nine FPs for the Pan-cancer cohort while the Pancancer* reported no FPs. This pattern was also evident across individual cancer types: without considering intracorrelation, FPs were reported in 8 cancer types, whereas with intracorrelation, FPs were observed only for Lymph-BNHL (Supplementary Data 7).

When we did not use an integrative approach and considered only recurrence or FI, the performance declined. In the FI-only setting, although no false positives were reported in either the pan-cancer or cancer-specific cohorts, the identified candidate drivers were limited to a few well-known cancer genes. In the recurrence-only setting, we observed more false positives compared to the original configuration, along with an overall decrease in iDriver performance. Notably, iDriver identified 78 significant hits that were missed by the recurrence-only setting, 48 of which were previously reported as cancer drivers in at least one of the CGC, OncoKB, or PCAWG gene lists. These included the CDS of *MED12* in Prost-AdenoCA, a known tumor suppressor in uterine leiomyoma and breast cancer^38^; the 3′UTR of *EPHA7* in Skin-Melanoma, whose CDS is a known driver in colorectal and melanoma cancers; and 3′UTR of *SETBP1* in Eso-AdenoCa, whose CDS is implicated as an oncogene or involved in fusion events in hematological malignancies—none of which were identified as cancer drivers in the PCAWG study. In addition, novel notable candidates missed by the recurrence-only method included the CDS of *FSTL5* in CNS-Oligo, *USP13* in Eso-AdenoCA, and the 3′UTR of *CNOT2* in CNS-PiloAstro, all of which are supported by existing literature (Table 1)^39–42^. Furthermore, iDriver, with its combined approach, can reduce the number of false positives typically reported by recurrence-based methods alone. For instance, the promoter of *PLEKHS1* is among the significant hits identified using recurrence-based iDriver in Bladder-TCC (q-value = 1.8e-5), Breast-AdenoCA (q-value = 0.011), Pan-cancer (q-value = 1.6e-6), and Pancancer* (q-value = 5.7e-9). However, it does not reach statistical significance when using the original iDriver. In the PCAWG study for cancer driver identification, the promoter of *PLEKHS1* was excluded from the driver list due to the presence of palindromic sequences that can be targeted by APOBEC^9^. The results of the ablation study under the different settings—excluding the uncorrelated setting due to its high false-positive rate—are presented in Figure 6bc.

### Functional and Literature Support for Candidate Drivers

In this study we introduced candidate drivers in coding and non-coding regions that were not reported in previous cancer gene lists. Table 1 provides a summary of the literature support for some of these genes and we provided a brief summary for some of them in this section. The first Pancancer* candidate not reported in the cancer gene lists is *EFNA1* core promoter. *EFNA1* is a member of the Ephrin family that are ligands of the largest family of the receptor protein-tyrosine kinases. Recent studies highlight *EFNA1* as a potent oncogene in cervical cancer, driven by a tumor-specific super-enhancer that amplifies its expression^43^. Mechanistically, *EFNA1* is transcriptionally regulated by FOSL2 and activates the EphA2-Src/AKT/STAT3 signaling axis, promoting tumor progression. Functional assays demonstrated that *EFNA1* knockdown in cervical cancer cell lines markedly inhibited proliferation, migration, invasion, and induced apoptosis and cell cycle arrest, while xenograft mouse models confirmed its essential role in tumor growth^43^. Furthermore, the *EFNA1* overexpression promotes gastric cancer liver metastasis by facilitating hepatic premetastatic niche formation through stimulating CCL2^44^. Another study demonstrates that Ephrin A1 functions as a ligand for EGFR, driving its activation, inducing epithelial-mesenchymal transition (EMT), and promoting the metastatic spread of gastric cancer cells^45^. Additionally, *EFNA1* is involved in the PI3K-Akt and MAPK signaling pathways and serves as a prognostic marker (Figure 4c). Other Pancancer* non-coding candidates that serve as prognostic cancer markers based on their RNA expression include *COL4A2*, *ARPC2*, *ISL1*, and *FOXG1* (Figure 4c).

*FOXG1* enhancer is another significant hit which is involved in the FoxO signaling pathway (hsa04068), which are transcription factors regulating cellular events like apoptosis and cell cycle. *FOXG1* is upregulated in lung cancer and promotes tumor cell proliferation by regulating the PI3K/AKT/mTOR pathway, as demonstrated by in vitro assays (MTT, CCK-8, colony formation, flow cytometry) and in vivo xenograft models^46^. Similarly, in glioblastoma, *FOXG1* overexpression cooperates with Wnt/β-catenin signaling to drive cell cycle re-entry from quiescence, with in vivo models confirming its role in gliomagenesis^47^. *PAF1* and *HNRNPK* are examples of the coding Pancancer* candidate drivers. In cervical cancer, *PAF1* knockdown reduces proliferation in vitro and slows tumor growth in vivo through inhibiting FLOT2-mediated MEK/ERK1/2 pathway^48^. Another study showed that PAF1 cooperates with YAP1 to drive acinar-to-ductal metaplasia and pancreatic ductal adenocarcinoma progression by regulating *SOX9* expression. In vitro and ex vivo studies demonstrated that *PAF1* knockdown or inhibition with CA3 and verteporfin suppresses the PAF1/YAP1/SOX9 axis, reducing pancreatic cancer cell growth, stemness, and duct-like structure formation^49^. Several studies have demonstrated the role of HNRNPK in cancer, with some showing that it directly regulates p53 and c-MYC and acts as a tumor suppressor, while others suggest it has an oncogenic role^50,51^. A more recent study demonstrated that overexpression of *HNRNPK* in human lung cancer cell lines promotes cell proliferation and enhances cell invasion, whereas *HNRNPK* knockdown suppresses cell proliferation and inhibits cell invasion^52^.

*ZIC1* CDS was identified as a novel candidate driver exclusively by iDriver in the Pancancer* cohort and in CNS-Medullo, whereas the population-level, Only-recurrence, and Only-FI methods failed to detect it as a significant hit (Figures 5a and 6b). In the CNS-Medullo cohort, *ZIC1* CDS harbored three mutations (two SNVs and one indel). Functional assessment of the SNVs using the AlphaMissense pathogenicity score indicated that both were pathogenic (Figure 5b)^53^. Several studies also support its role in cancer^54–56^. Specifically, a recent study showed that *ZIC1* functions as a context-dependent cancer driver in medulloblastoma, exhibiting loss-of-function alterations in Group 4 tumors and gain-of-function mutations in SHH subtypes. Functional assays reveal that *ZIC1* overexpression suppresses Group 3 tumor growth but promotes SHH medulloblastoma proliferation, underscoring its dual oncogenic and tumor-suppressive roles in medulloblastoma^36^. *FSTL5* CDS, a candidate called by iDriver for CNS-Oligo, is a prognostic marker in Glioma (Figure 5c). *FSTL5* downregulation in hepatocellular carcinoma correlates with poor prognosis. In vitro and in vivo studies show that *FSTL5* overexpression inhibits hepatocellular carcinoma growth by promoting caspase-dependent apoptosis through regulation of Bcl-2 family proteins^57^, and support its tumor suppressive role in hepatocellular carcinoma^39^. Another example of the cancer-specific CDS candidate of our study, with several literature supports across multiple tumor types, is *USP13* in Eso-AdenoCa cohort. In gastric cancer, USP13 deubiquitinates and stabilizes cyclin D1, enhancing cell cycle progression and tumor growth both in vitro and in vivo.^40^. In *KRAS*-mutant non-small cell lung cancer, USP13 removes K63-linked ubiquitin from β-catenin, enhancing β-catenin and TCF4 binding, resulting in promoting metastasis and poor prognosis^41^. In clear cell renal cell carcinoma, USP13 stabilizes ZHX2, which promotes tumor cell proliferation and invasion through MEK/ERK signaling activation. In vivo experiments further demonstrate that USP13 is essential for tumor growth^58,59^. Furthermore, in breast cancer, USP13 promotes metastasis by regulating Twist1 stability via a negative feedback loop between USP13 and Twist1^60^.

An example of iDriver non-coding candidates in cancer-specific cohorts is *PAX9* 5’UTR in Eso-AdenoCa. Evidence supports *PAX9* as a tumor suppressor gene across multiple epithelial cancers. In cervical cancer, in vitro and in vivo experiments show that *PAX9* overexpression inhibits cell proliferation and promotes apoptosis^61^. Similarly, in oral squamous cell carcinoma, pharmacological reactivation of silenced *PAX9*, a gene involved in differentiation of oral squamous cell carcinoma, through DNA methyltransferase inhibition restores its antitumor function, further validating its role in suppressing malignant progression^62^. Further candidates introduced by iDriver with their support are summarized in Table 1.

## Discussion

In this study, we introduce iDriver, a probabilistic framework for the discovery of cancer driver elements across both coding and non-coding genomic regions. By incorporating patient-specific mutational heterogeneity, functional impact scores, and BMR modeling informed by 1,500 (epi)genomic features, iDriver provides a unified strategy to detect signals of positive selection. Our benchmarking analyses demonstrate that iDriver consistently outperforms state-of-the-art driver discovery algorithms—both recurrence-based (e.g., MutSig, LARVA, NBR) and functional impact-based approaches (e.g., OncodriveFML)—across 31 cancer cohorts, encompassing both pan-cancer and tissue-specific analyses.

Unlike conventional methods that do not account for differences in mutational burden across patients, iDriver explicitly corrects for inter-patient heterogeneity in mutation rates—an important yet overlooked confounder. By integrating patient-level functional impact distributions and employing a Poisson–Binomial framework to model heterogeneous mutation probabilities, iDriver ensures that mutations in hypermutated samples contribute proportionally less to driver inference. This adjustment increases the number of significant discoveries while maintaining stringent control of false positives.

Applying iDriver to the pan-cancer and cancer-specific cohorts revealed both coding and non-coding elements. Notably, iDriver uncovered previously unreported candidate drivers such as *EFNA1*, *COL4A2*, *ARPC2*, and *FOXG1*, several of which show strong prognostic associations across multiple tumor types. Independent in vivo and in vitro studies support their mechanistic involvement in oncogenic signaling pathways, including PI3K–AKT, MAPK, and Wnt. The discovery of tissue-specific drivers—such as *RYR2* CDS in CNS-GBM, participating in calcium signaling, and *RPS6KA3* CDS in liver HCC within the MAPK pathway, despite being absent from COSMIC CGC and OncoKB—further demonstrates iDriver’s capacity to capture context-dependent selective pressures.

A key strength of iDriver lies in its multilevel modeling of heterogeneity: (i) across cohorts, through cohort-specific BMR estimation and driver discovery; (ii) across genomic regions, by integrating (epi)genomic features such as replication timing and conservation; and (iii) across patients, by accounting for individual mutation burdens and functional impact distributions.

Our ablation analyses highlight the necessity of this integrative design. Removing patient-specific burden correction or ignoring mutation correlation within elements markedly reduced sensitivity and increased false positives, respectively. In contrast, the complete iDriver configuration identified biologically meaningful drivers without inflation of type I error. Importantly, incorporating intra-element correlation through a mixed-effects model substantially reduced spurious hits observed in earlier burden-based methods, emphasizing the importance of modeling both the spatial clustering and functional coherence of mutations.

From a biological perspective, the novel drivers identified by iDriver provide mechanistic and translational insights. For example, *ZIC1* in CNS-Medullo and *FSTL5* in CNS-GBM—supported by independent evidence and/or survival associations—highlight iDriver’s capacity to uncover clinically relevant targets that may be missed by recurrence-only or population-level approaches.

Overall, by incorporating individual mutation burdens and functional impact, iDriver achieves robust driver prioritization even in hypermutated samples. The integration of CADD-based functional scores with recurrence data maximizes detection power while minimizing false positives. The probabilistic graphical model, with parameter approximation via a Beta distribution and using a mixed-effect model for significance testing, provides a rigorous statistical foundation applicable to both coding and non-coding elements. Benchmarking analyses demonstrate iDriver’s superior performance relative to existing methods, with high specificity and biological interpretability. Mapping the identified drivers to signaling pathways and survival outcomes further underscores their translational relevance in oncology.

Despite its strengths, iDriver has several limitations. The current implementation uses CADD scores, which may not fully capture context-dependent or structural effects of mutations. Moreover, while iDriver models SNVs, indels, and MNVs, it does not yet incorporate structural variants, copy-number alterations, or epigenetic driver events, which also contribute to oncogenesis. Expanding iDriver to integrate these additional layers of variation represents an important direction for future development.

Collectively, iDriver provides a robust and statistically principled framework for pan-cancer driver discovery, combining patient-specific mutational heterogeneity with functional impact scores to link statistical rigor to biological insight. The resulting catalog of drivers provides a valuable resource for further exploration and the identification of novel therapeutic targets across diverse cancer types.

## Methods

### Whole-genome somatic mutation data

We used whole-genome somatic SNVs and indels from 2,583 white-listed PCAWG donors, obtained from the ICGC Data Portal (dbGaP Project ID for the TCGA portion: 32607, accession: phs000178), as part of the ICGC 25k Release. We focused on 29 cohorts with at least 18 donors each, as well as two Pan-cancer cohorts. The primary Pan-cancer cohort includes 2,514 non-hypermutated donors. Donors with more than 30 mutations per Mb were categorized as hypermutated donors. The Pancancer* cohort is identical to the Pan-cancer cohort but excludes samples from the Skin-Melanoma, Lymph-CLL, Lymph-NOS, and Lymph-BNHL cohorts. The main analyses were conducted on non-hypermutated samples. However, we also analyzed cohorts that included both hypermutated and non-hypermutated samples, as well as a separate Pan-cancer cohort (n=69) consisting exclusively of hypermutated donors (the Hypermutated cohort).

### Simulated neutral somatic mutation dataset

We used the Sanger-simulated data from PCAWG to assess the type I error of driver discovery methods. This dataset of neutral somatic mutations captures the heterogeneity of BMRs across cancer types and among different patients within each type, while preserving the mutational signatures (see Rheinbay et al. for more details ^9^).

### Genomic and epigenomic predictors of mutation rate variation

We used 1,500 genomic and epigenomic features known to influence the BMR. These features are broadly categorized into nine groups. Seven of these groups (n = 1372) were previously used in DriverPower and eMET and are described in those studies^24,28^. Briefly, the conservation group includes three features derived from phastCons^63^ and phyloP scores^64^. The Replication timing group consists of 16 features based on Repliseq processed signals from different cell types of origin. The nucleotide content group comprises the frequencies of 46 di- and tri-nucleotide sequences within each element of interest. The epigenetic marks group contains 979 features, including various histone methylation and acetylation marks from multiple cell types obtained from the Roadmap Epigenomics consolidated data^65^. The RNA expression category includes 127 gene expression features also from the same source. The chromatin compartments group consists of 12 features derived from the HiC data across different normal or cancer cell lines, encoded as active/inactive compartments in BigWig-format file. The DNA accessibility group includes 189 features including open chromatin data from the ENCODE project and MACS2 DNase narrow peak signals from Roadmap Epigenomics data, covering various normal and cancer cell lines.

In addition to these, we incorporated two new sets of features (Supplementary Data 8): 1) genomic positions of DNA secondary structures targeted by APOBEC3A, as previously used in MutSpot^26^. 2) 127 DNA methylation features from various normal and cancer cell lines, based on Roadmap Epigenomics consolidated data, since methylated regions are known to exhibit elevated mutation rates^66^.

### Scoring the deleteriousness of somatic mutations with CADD

We used CADD scores (GRCh37-v1.6) to assess the deleteriousness of somatic mutations—including SNVs, MNVs, and indels—identified in the cohorts^29^. CADD integrates annotations from evolutionary, functional, regulatory, and structural sources, providing broad scores across both coding and non-coding regions and across different mutation types. A higher CADD score indicates a greater likelihood of deleterious impact. Pre-scored files were used for SNVs and indels, while MNVs and a subset of indels without precomputed scores were annotated using the CADD web interface.

### Population-level BMR estimation

To estimate the BMR of genomic elements, we used eMET that models background mutation rate as a function of genomic features. Definitions of the genomic elements were adopted from the original eMET study. We applied eMET independently to each patient cohort to obtain population-level mutation rates. For each element *e*, eMET outputs the expected mutation rate of the element *e* under neutral mutational processes in a cohort of *N* patients.

### iDriver algorithm

The iDriver algorithm is applied independently to each cancer cohort. Below, we outline the algorithmic steps for a given genomic element of interest, denoted as *e*. For clarity, we omit *e* from the notation when it is obvious from the context. Assume there are *N* patients in the cohort. The algorithm proceeds as follows:

#### Population-level Background Mutation Rate Estimation

We first obtained the BMR for the genomic element *e* using eMET, denoted as *p_BMR_*. Since *p_BMR_* is normalized by both the number of patients in the cohort, *N*, and the length of the genomic element in base pairs, *L*, in the original publication, we transform it into the expected total number of predicted mutations in the element:

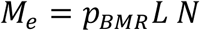

The probability that element *e* is mutated in a single patient, denoted as *p_e_*, is then computed by scaling to the observed cohort-wide mutation burden:

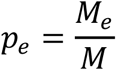

where *M* is the total number of observed mutations in the cohort.

#### Patient-specific signals of positive selection

To quantify the signals of positive selection at the individual-patient level, we defined a score *t_i_* for each patient *i*. Patients without mutations in element *e* were assigned *t_i_* = *0*, indicating no evidence of positive selection. For patients harboring *n* mutations in element *e*, each with functional scores *s_1_*, . . . , *s_n_* , we calculated the observed total score as *Σ_obs_* = *s_1_*+. . . +*s_n_*. We then computed the probability that the sum of mutation scores under the background model is less than the observed sum, i.e., *t_i_* = *P*(*Σ* < *Σ_obs_*), where *Σ* represents the sum of mutation scores under the null model. Larger values of *t_i_* indicate stronger evidence that the observed signal deviates from the background distribution, thus suggesting positive selection in patient *i*. To estimate *t_i_*, we computed:

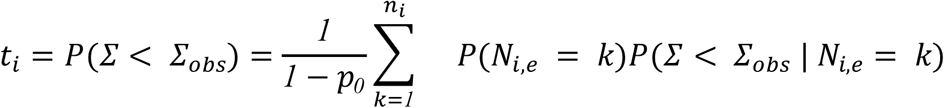

where *N_i_*_,*e*_ is the number of mutations expected in patient *i* for element *e* under the null (background) model, assumed to follow a binomial distribution *N_i_*_,*e*_ ∼ *Bin*(*n_i_*, *p_e_*). Here, *n_i_* is the total number of mutations in patient *i* and *p_e_* the background probability of observing a mutation in element *e*. The normalization factor (*1* − *p_0_*) accounts for the fact that the summation starts at *k* = *1*, where *p_0_* = *P*(*N_i_*_,*e*_ = *0*) is the probability mass at zero.

The conditional distribution of total scores given *N_i_*_,*e*_ = *k* was modeled as a normal distribution with mean *kμ* and variance *kσ* ^2^, i.e., *Σ* | *N_i_* = *k* ∼ *N*(*kμ*, *kσ* ^2^), reflecting the assumption that total summation of mutation scores scale linearly with the number of mutations. For computational efficiency, the summation over *k* was truncated at a maximum value *C*. Specifically, we approximated

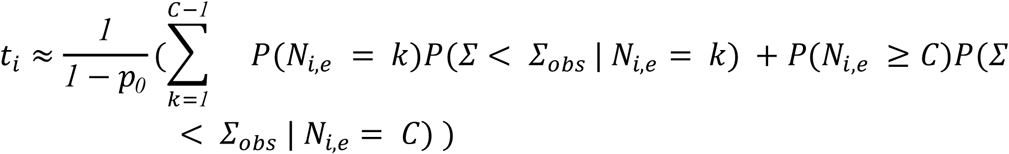

In practice, we set *C* = *4*, which provides a good approximation since the probability of larger counts is negligible and dominated by the term *P*(*N_i_*_,*e*_ = *C*).

#### Estimation of μ and σ

The parameters μ and σ were estimated from observed functional scores of mutations in putative non-driver elements. Estimation was performed separately for each element type and cohort to account for context-specific variability. The resulting values of μ and σ are provided in Supplementary Data 9.

#### Statistical testing for positive selection at the element level

To test whether a genomic element *e* is under positive selection, we use the aggregated patient-level scores as the test statistic. Specifically, we define the observed statistic as

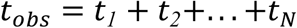

where *t_i_* is the patient-specific positive selection score (defined earlier) and *N* is the number of patients in the cohort. Under the null hypothesis (no selection), the corresponding null distribution is

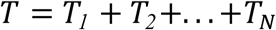

where each *T_i_* represents patient-specific positive selection scores under the null assumption. Since *T_i_* takes value zero (if non-mutated) or a nonzero score, we improved the approximation by marginalizing *P*(*T* > *t_obs_*) with respect to the random variable *N_nz_*, the number of patients harboring at least one mutation in element *e* under the null. Thus, the p-value for element *e* is computed as

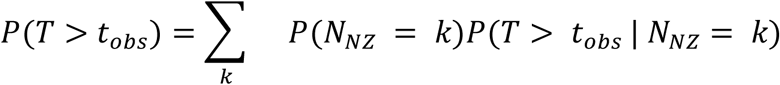

The distribution *P*(*N_NZ_* = *k*) was obtained from the Poisson–Binomial distribution. For each patient *i* ∈ {*1*, . . . , *N*}, the probability of having at least one mutation in element *e* was defined as

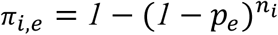

The set of probabilities {*π_1_*_,*e*_, *π_2_*_,*e*_, . . . , *π_N_*_,*e*_} defines the Poisson–Binomial distribution from which *P*(*N_NZ_* = *k*) is computed. Accounting for heterogeneous mutation probabilities across patients provides a rigorous null model and ensures proper type I error control.

#### Normal approximation for Beta-sum tail probability

The conditional term *P*(*T* > *t_obs_* | *N_NZ_* = *k*) was computed under the assumption that each *T_i_* follows a Beta distribution, *T_i_* ∼ *Beta*(*α*, *β*). By the central limit theorem, the summation of these terms can be approximated by a normal distribution, *T* ∼ *Normal*(*μ_T_*, *σ^2^_T_*) where

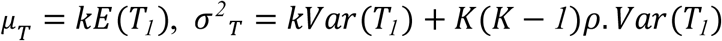

and ρ is the intra-class correlation reflecting that some genomic regions may harbor variants with correlated CADD scores. For the Beta distribution, we have

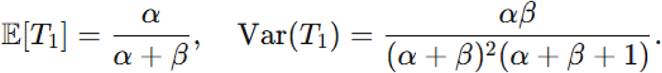

Since the *T_i_* are identically distributed, the above moments are derived with respect to *T_1_* without loss of generality. The tail probability is then approximated as

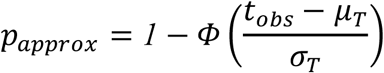

where *Φ* is the cumulative distribution function of the normal distribution.

#### Intra-class correlation (ρ) estimation

To account for correlation among mutation scores within the same genomic element, we estimated the intra-class correlation coefficient (*ρ*) using a linear mixed-effects model. Specifically, for each pan-cancer cohort, we selected patient-level scores *t_i_* from genomic elements with non-zero within-element variance. We fit a random-intercept model of the form (using the *lme* function from the *nlme* package in R):

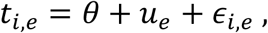

where *θ* the overall mean, *u_e_* ∼ *N*(*0*, *σ^2^_between_*) is the random intercept for element e, and *ε_i_*_,*e*_ ∼ *N*(*0*, *σ^2^_witℎin_*) is the residual variance. From the fitted model, we extracted the variance components and computed *ρ* as:

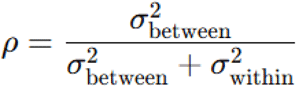

Because intra-element correlation varies with element length, we stratified genomic elements of each element type (e.g. CDS, 3’UTRs) into four length-based categories (defined by quantiles of element lengths for each element type) and computed a distinct ρ for each category.

#### Beta distribution parameter estimation

To estimate the Beta distribution parameters *α* and *β*, we applied the method of moments using the same set of observed patient-level scores (*t_i_*) and categories as what we used for estimating ρ. Let

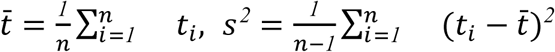

denote the sample mean and variance of the observed patient-level *t_i_* scores, computed separately for each category with *n* scores. The parameters are then estimated as:

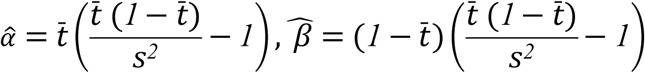

We used the estimated parameters α, β, and ρ from the Pancancer cohort, which included mutations from both hypermutated and non-hypermutated donors, for all cancer-specific cohort analyses. The estimated values of *α*, β, and *ρ* for the Pancancer cohort are provided in the Supplementary Data 10.

### Multiple-testing correction

Multiple-testing correction was performed using the Benjamini–Hochberg procedure, applied separately to coding and non-coding elements that harbored at least two mutations within each cohort. Elements with a q-value < 0.1 were considered statistically significant candidate drivers.

### Gold Standard

To define the list of known cancer genes, we used the COSMIC Cancer Gene Census (CGC) v101^32,38^, the OncoKB cancer gene list (April 2023)^67,68^, and the union of 603 CGC v80 genes with 369 genes (n = 757) identified by Martincorena et al. and Lawrence et al. in their WXS studies^19,69^, as provided in Supplementary Table 7 of the Rheinbay et al. study^9^. We also included all PCAWG IDs reported as cancer drivers by PDFIWG in Supplementary Tables 4 and 5 of the Rheinbay et al. study, with a “Pre-filter q-value” ≤ 0.1 (referred to as PCAWG (raw)). For the analysis of the Hypermutated cohort, we further defined a category termed PCAWG (post-filtering), which excluded PCAWG (raw) elements that were not present in CGC or OncoKB, and did not meet the Post-filter q-value ≤ 0.1 criterion. Throughout the paper, the term PCAWG refers to the PCAWG (raw) elements.

We defined the CGC gene list as our gold standard, as it is a curated collection supported by strong evidence of involvement in cancer development. When benchmarking iDriver against other driver discovery methods, we defined true positives as follows: For CDSs, 5’UTRs, promoters, and 3’UTRs, significant hits (q-value < 0.1) were considered true positives if they overlapped with genes listed in the CGC. For enhancers, a significant hit was classified as a true positive if any of its predicted target genes—based on gene-enhancer mapping provided by PCAWG—appeared in the CGC cancer gene list.

### Ablation study

In addition to the original iDriver, which incorporates functional impact and mutation recurrence, and accounts for the correlation of scores among elements of the same length category, we performed a series of ablation experiments. In each experiment, one component was removed to assess its influence on driver discovery across cancer cohorts. For the population-level setting, we eliminated the effect of mutational burden heterogeneity among patients within each cohort by replacing individual burdens with the average mutational burden across all patients, ensuring that all samples shared the same *n_i_*. All other algorithmic steps remained identical to those used in the original iDriver analysis. The uncorrelated score setting was performed in the same manner as the original analysis, except that ρ was set to 0.

For the recurrence only setting (FI-free setting), the observed statistic was calculated as the probability that the sum of mutations under the background model is less than the observed number of mutations, without the conditional normal distribution of total scores given *N_i_*_,*e*_ = *k* term that was used in the original setting, and all other algorithmic steps remained identical to those used in the original iDriver analysis.

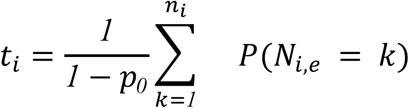

For the FI only setting (Burden-free setting), the only difference with the original setting is in the calculation of the patient-specific positive selection scores under the null assumption, where the probability of having at least one mutation in element *e* in respect to the mutational burden of patients is not considered, and each *T_i_* follows a Beta distribution, *T_i_* ∼ *Beta*(*α*, *β*). therefore it is computed as

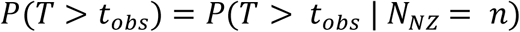

Where *n* is the number of mutations in element *e*.

### Pathway information

Information on signaling pathways associated with the significant hits was obtained from the Kyoto Encyclopedia of Genes and Genomes (KEGG) PATHWAY database (https://www.genome.jp/kegg/). For each significant gene symbol, the corresponding Entrez ID was retrieved using the mapIds function from the *AnnotationDbi* R package. Subsequently, KEGG pathways corresponding to each Entrez ID were obtained using the *keggLink* function from the *KEGGREST* package. The relevant KEGG networks and network elements related to cancer or infection-associated pathways were retrieved through the *KEGGREST* package by specifying human (“hsa”) KEGG pathway identifiers. Using the “nt” network identifiers for each hsa pathway, we extracted the list of genes included in each pathway. Detailed information on network elements involved in cancer-related pathways, along with their corresponding gene lists, is provided in Supplementary Data 11. For the Pathways in Cancer (hsa05200), we used data provided in Table S2 of the Mina et al. study^70^.

### Analysis by mutation type

In addition to pooling all mutations for the original iDriver implementation, we also applied iDriver separately to different mutation types. In one setting, iDriver was run independently on synonymous and nonsynonymous mutations. To achieve this, all variants located within CDSs were annotated using FunSeq2^31^. Variants annotated as “nonsynonymous”, “prematureStop”, “removedStop”, or “spliceOverlap” were grouped as nonsynonymous, while variants annotated as “synonymous” were grouped as synonymous.

For each group, we applied the element-specific model described in the eMET study^28^ across different cancer types and estimated the number of mutations for each element and cohort and calculated pₑ. For each cohort, we then computed the mean and variance of CADD scores separately for the synonymous and nonsynonymous groups, and subsequently implemented iDriver on each group independently.

We also performed separate modeling of SNVs and non-SNVs. To this end, we estimated the number of SNV and non-SNV variants using distinct eMET models for each variant group across cancer cohorts, followed by the calculation of pₑ and the mean and variance of CADD scores separately for SNVs and non-SNVs. Finally, iDriver was applied using the original setting.

### Combining p-values

We used Fisher’s method and Tippett’s method to integrate evidence of positive selection obtained from applying iDriver separately to SNVs and non-SNVs, using the *poolr* package in R. Briefly, for each element, if a p-value was available for only one variant type (SNV or non-SNV), that p-value was reported. For elements with p-values available for both variant types, a combined p-value was calculated.

Fisher’s method combines the two p-values using a test statistic that follows a chi-square distribution with two degrees of freedom. In contrast, Tippett’s method takes the smallest of the two p-values as the test statistic and evaluates its significance accordingly.

### Type I error analysis

We applied iDriver to simulated data to assess the Type I error rate. After performing multiple-testing correction on all tests—either jointly for all coding and non-coding elements or separately for coding and non-coding elements with at least two mutations—we reported the number of significant hits with a q-value < 0.1, as determined using the Benjamini–Hochberg procedure.

### Analysis of hypermutated donors

To apply iDriver to hypermutated donors, we estimated the predicted number of mutations using three different approaches: i) by directly estimating mutation counts from the hypermutated cohort itself using eMET, ii) by applying eMET to the Pancancer cohort and subsequently adjusting the predicted mutation counts based on the ratio of mutation rates between the hypermutated and Pan-cancer cohorts, and iii) by applying eMET to the Pancancer* cohort and adjusting the predicted mutation counts using the corresponding ratio between the hypermutated and Pancancer* cohorts. We further defined a local BMR for each element, calculated from the mutation rate within a ±50 kb window around the element, as described in a previous study^28^. Elements exhibiting estimated local mutation rates exceeding 5×, 10×, 15×, or 20× the estimated rates using the genomic and epigenomic features, were filtered out in subsequent analyses.

We then compared iDriver performance across the different estimation and filtering strategies and against five random subsamples of non-hypermutated donors matched to the hypermutated cohort in cohort composition (number of donors per cancer type). The number of patients for each cancer type and the total number of mutations in the hypermutated cohort and matched subsamples are provided in Supplementary Data 12. For this analysis, the FDR was recalculated after filtering the PCAWG (raw) elements, separately for coding and non-coding elements with at least two mutations.

### Data availability

All of the datasets used in this study are publicly available and listed in Supplementary Data 8 (feature URLs) and Supplementary Table 1. The TCGA somatic SNV/indel data, and the TCGA portion of the Sanger simulated mutations with controlled access policies were obtained from dbGaP (Project ID: 32607; accession: phs000178).

### Code Availability

All source codes used for implementing iDriver are publicly available on GitHub at git clone https://github.com/FaridehBahari/iDriver.git.

## Acknowledgments

This work is based upon research funded by Iran National Science Foundation (INSF) under project No. 4022073. We sincerely thank Dr. Mina Karimpour for the insightful discussions and valuable suggestions. We accessed the TCGA portion of somatic mutation calls of the PCAWG project through dbGaP (Project ID: 32607, accession: phs000178).

## Authors’ contributions

Both authors conceptualized and planned the project, and developed the methodology. F.B. implemented the method in R and conducted all the experiments and visualizations. H.M. supervised the study and verified the findings of this work. Both authors wrote the manuscript and agreed to the final version of the manuscript.

