## Supplementary Information for "iDriver: A patient-centric framework for genome-wide cancer driver discovery"

### Cancer Driver Gene Discovery: A Patient-Level Statistical Framework

#### Supplementary Note 1: Excluding CompositeDriver from benchmarking methods

One of the driver discovery methods used by PCAWG is CompositeDriver[1]. In this approach, the functional impact scores of all mutations within a genomic element are summed to obtain a score, which is then compared against a null distribution to calculate p-value. The null model is generated by randomly sampling the same number of mutations observed in the given element from all mutations of the same element type, matched for replication timing, nucleotide context, DNase I hypersensitive sites, and histone marks.

CompositeDriver relies on functional impact scores from FunSeq2[2]. FunSeq2 integrates multiple genomic and epigenomic features to prioritize variants, including prior knowledge of cancer driver genes, DNA repair genes, and actionable genes. Consequently, the use of such prior information introduces information leakage, potentially inflating performance when benchmarking driver discovery methods. Therefore, to ensure a fair and unbiased comparison, in our study we benchmarked iDriver against all PCAWG methods except CompositeDriver.

**Supplementary Note 2: Hypermutated cohort**

We defined a cohort consisting of all hypermutated cases (n = 69) and applied iDriver using three different approaches for background mutation rate ($p_e$) estimation, as described in the Methods section: (1) eMET-based estimation for the hypermutated cohort, (2) updated Pan-cancer estimation including hypermutated donors, and (3) updated Pancancer* estimation excluding hypermutated donors. We compared the performance of iDriver under these three $p_e$ estimation strategies, with and without additional filtering of elements exhibiting elevated local mutation rates (±50 kb window) that exceeded the predicted mutation rates from (epi)genomic features by 5×, 10×, 15×, or 20×.

Our analyses showed that using Pan-cancer or eMET-based estimates for $p_e$ yielded fewer significant hits compared to Pancancer* estimates, which resulted in a large number of significant elements at a false discovery rate (FDR) of 0.1 (n = 1,982 coding and n = 5,110 noncoding elements; Supplementary Data 1). To determine the optimal $p_e$ estimation strategy, we further examined the top 100 coding and noncoding elements identified by iDriver under each approach. This analysis revealed that employing updated Pan-cancer estimates without applying any local mutation rate filters provided the best performance for the hypermutated cohort, for both coding and noncoding elements (Supplementary Figure 7a).

Additionally, when comparing the number of CGC, OncoKB, or PCAWG (post-filter) genes among the top 50 iDriver hits for the hypermutated cohort with those from five subsampled non-hypermutated donor sets, we observed comparable numbers of known and novel elements across all six cohorts (Supplementary Figure 7b). These findings highlight the robustness of iDriver in identifying cancer-relevant elements, even when applied exclusively to a hypermutated cohort.

**Supplementary Note 3: Separate modeling of SNVs and non-SNVs for cancer gene prioritization**

Since SNVs, doublets, MNVs, and indels are not necessarily concordant and may arise from distinct mutagenic processes, we separately estimated the BMR for SNVs and non-SNVs and subsequently applied iDriver using these estimates. The resulting p-values were then combined using two approaches: Fisher's method and Tippett's method. Comparing these results with those obtained by pooling all variant types showed that the pooled approach (iDriver) outperformed both Fisher's and Tippett's combinations in most cohorts (Supplementary Figure 8). Specifically, iDriver achieved the best performance in 16 and 10 cohorts for coding and non-coding elements, respectively. Fisher's method performed best in four cohorts for both coding and non-coding elements, while Tippett's method performed best in three cohorts. For the remaining cohorts, at least two of the methods yielded comparable performance.

**Supplementary Note 4: Exploring the enrichment of synonymous mutations in coding elements**

We applied iDriver to multiple cohorts, each containing at least 20 donors with synonymous mutations. To this end, we restricted the analysis to synonymous variants within CDS regions. For each cohort, we computed the mean and variance of CADD scores across all synonymous mutations as inputs for iDriver. The expected number of mutations per CDS was estimated using cohort-specific, element-specific models[3].

In total, we identified 19 significant CDSs (q-value < 0.1), including 15 in the Lymph-BNHL cohort, nine in the Pan-cancer cohort, and two in the Eso-AdenoCa cohort (Supplementary Data 2). The majority of these hits correspond to known cancer genes listed in the CGC or OncoKB databases, including *BCL2*, *PIM1*, *MYC*, *SGK1*, *SOCS1*, *DTX1*, *BTG1*, *DUSP2*, *IRF4*, *CTNNA2*, and *TP53*. After post-filtering, only *TP53* remained significant in the PCAWG dataset (Supplementary Figure 9)[4], and *CTNNA2* was not among the PCAWG cancer gene list. This finding is consistent with previous studies reporting positive selection on certain synonymous mutations in *TP53*[5,6].

### Supplementary Figures

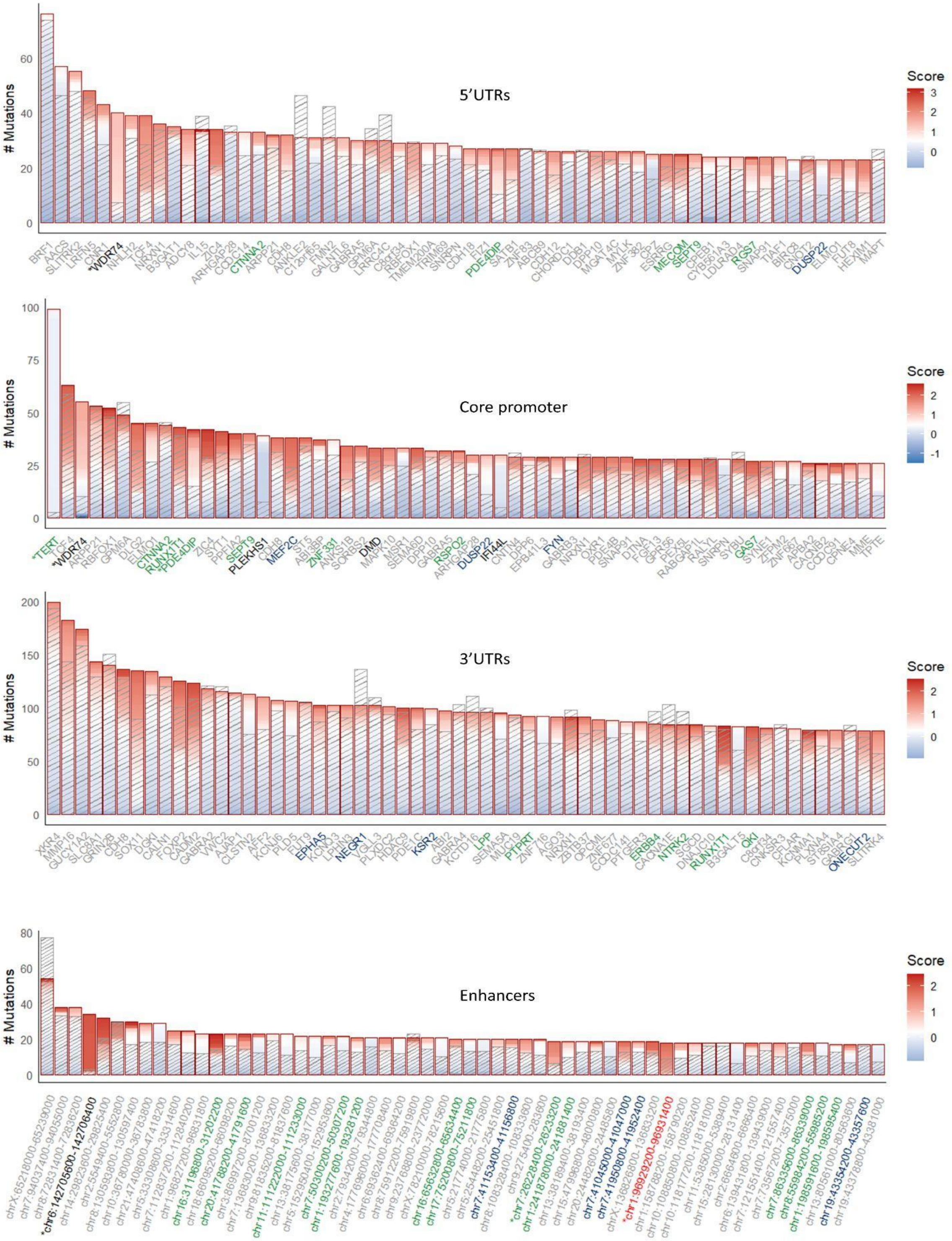


**Supplementary Figure 1** The top 60 most frequently mutated non-coding elements, showing both observed (highlighted by red boxes around the stacks) and expected numbers of mutations estimated using eMET (indicated by diagonal stripes in the bars). Mutations within each element are represented as stacked bars, colored according to their CADD pathogenicity scores, with the white color indicating the average of the scores in the corresponding element type. Gene symbol or enhancer coordinates colors indicate whether the corresponding gene is listed as a COSMIC CGC gene (green), an OncoKB cancer gene (blue), a PCAWG cancer gene (black), novel candidates called by eMET (red), or not listed in any cancer gene database (grey). Significant hits identified by iDriver are highlighted with an asterisk beside the gene name.

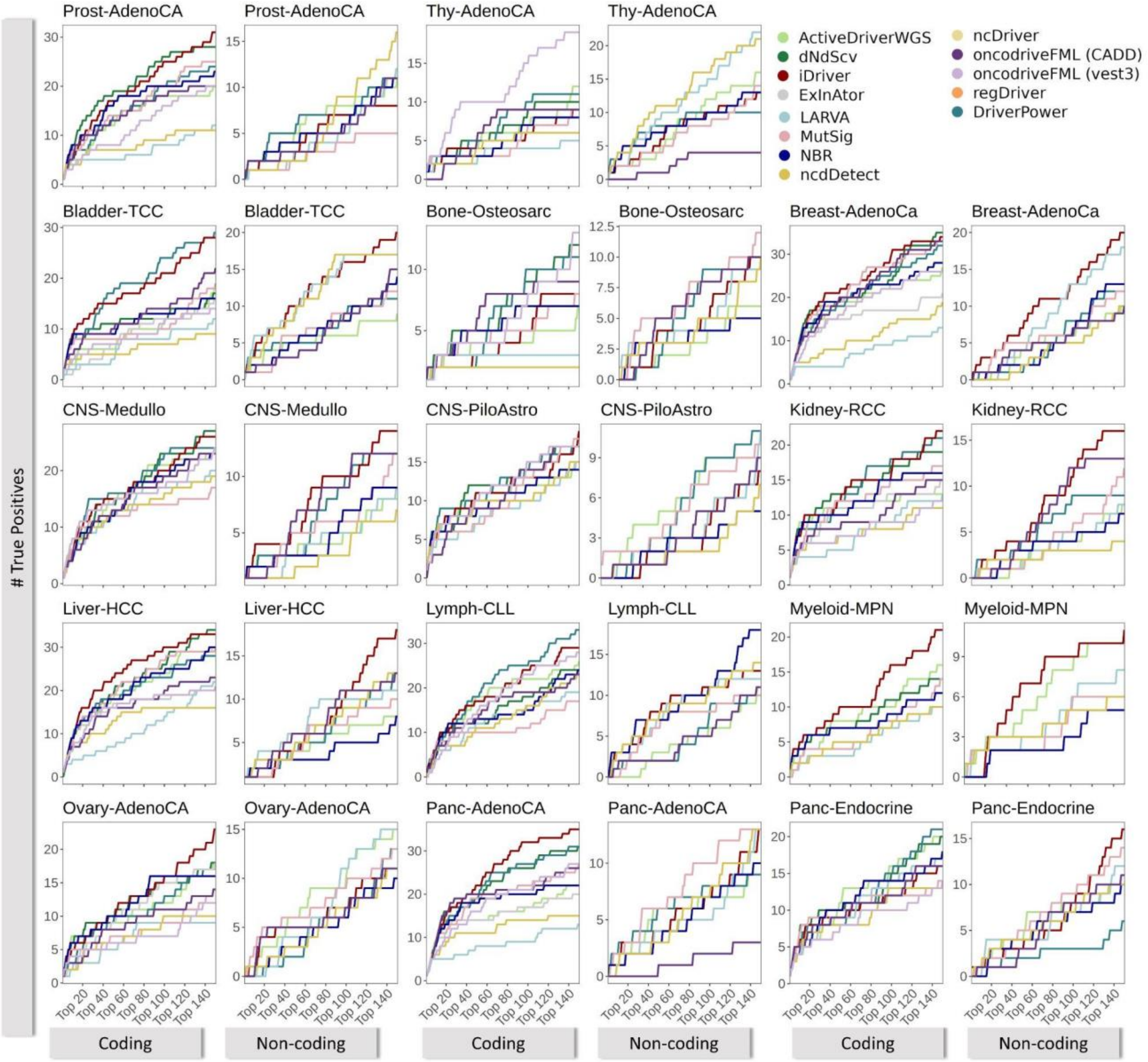


**Supplementary Figure 2** Cumulative number of TPs among the top-ranked elements (up to top 150) for different cancer types without any hypermutated donors for coding and non-coding elements compared to 12 driver discovery methods.

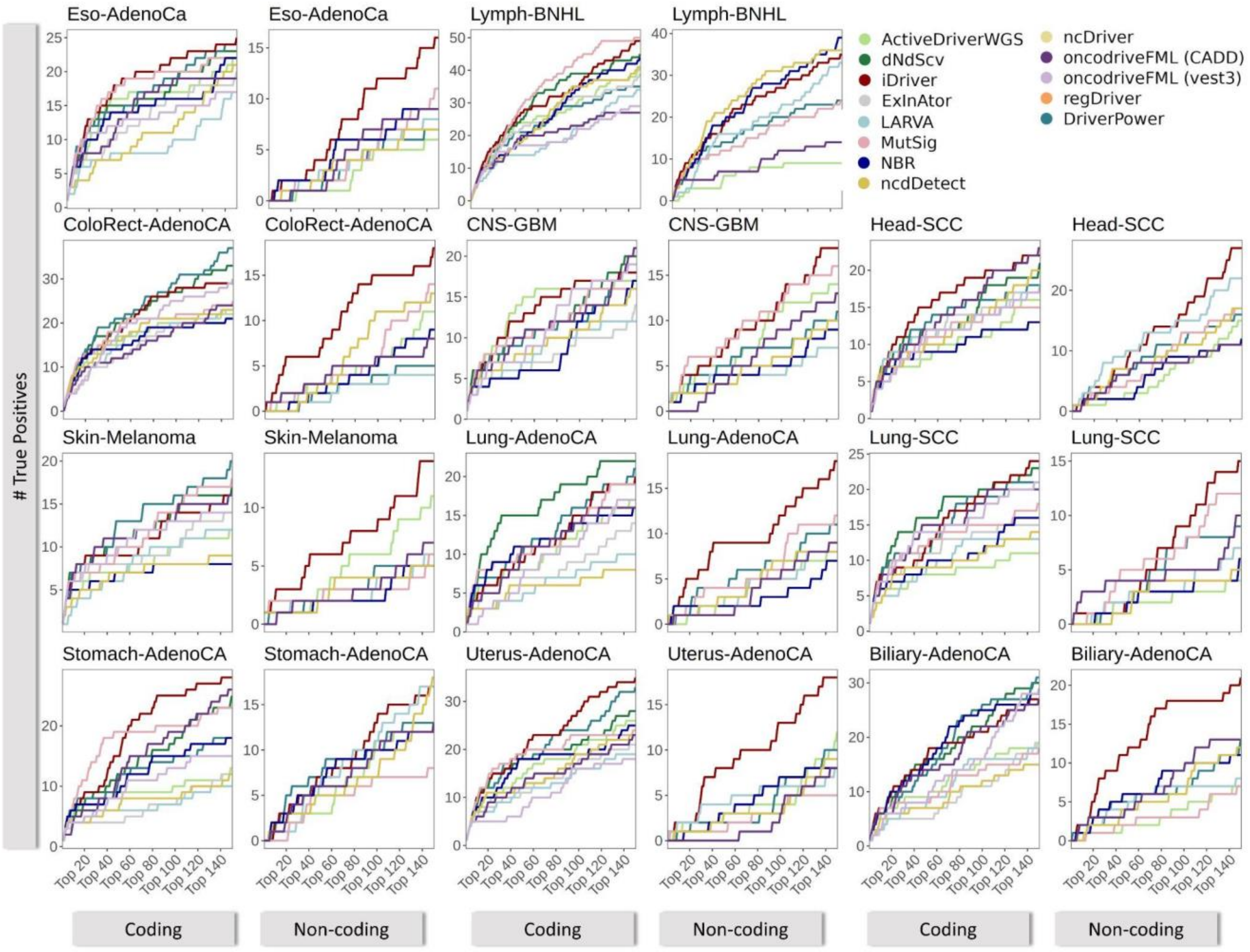


**Supplementary Figure 3** Cumulative number of TPs among the top-ranked elements (up to 150) across cancer types, excluding hypermutated donors, for coding and noncoding elements, compared with 12 driver discovery methods.

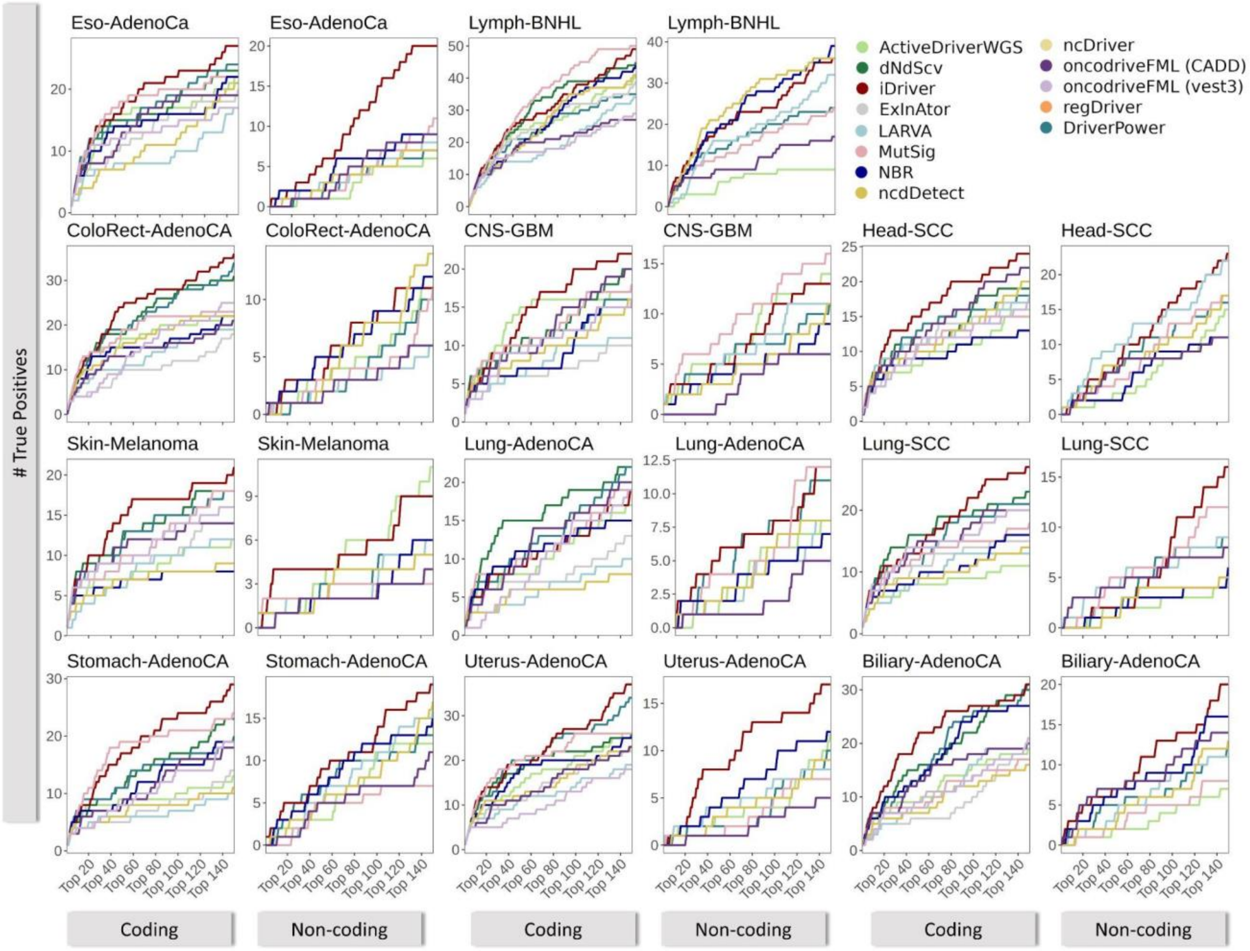


**Supplementary Figure 4** Cumulative number of true positives (TPs) among the top-ranked elements (up to 150) across cancer types, with all donors including hypermutated donors, for coding and noncoding elements, compared with 12 driver discovery methods.

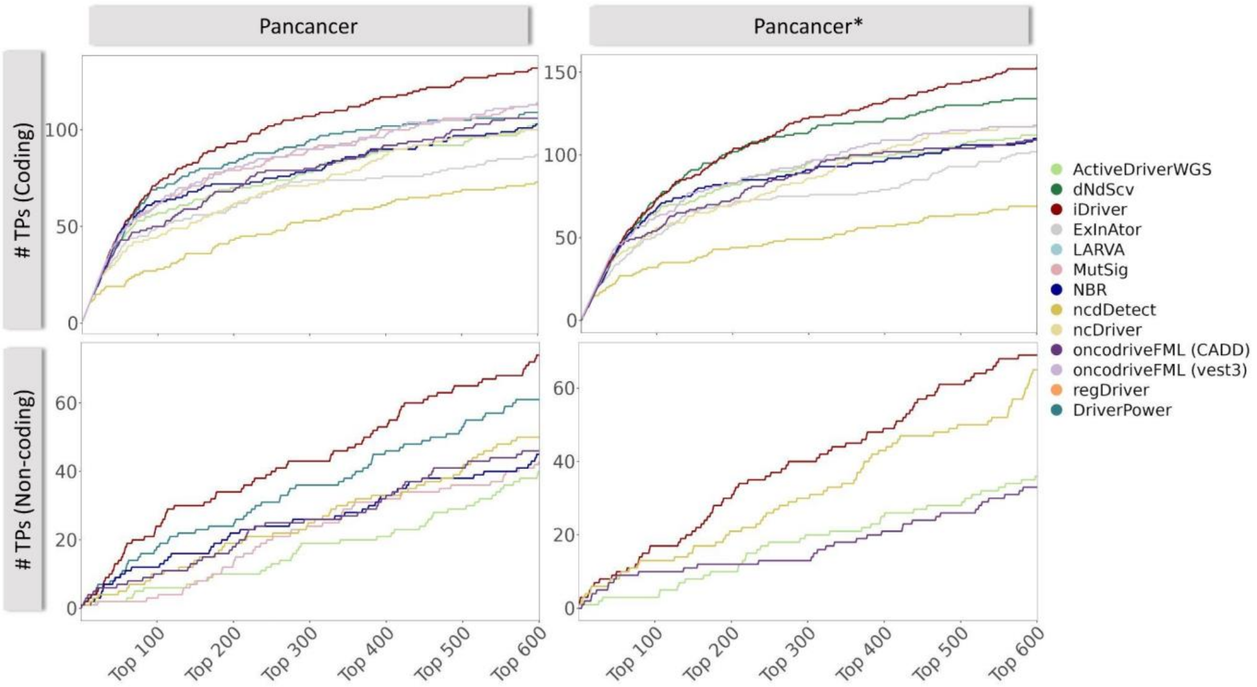


**Supplementary Figure 5** Cumulative number of true positives (TPs) among the top-ranked elements (up to top 600) for the Pancancer and Pancancer* cohorts without filtering for hypermutated donors compared to 12 driver discovery methods. These plots show the superior ranking ability of iDriver with all donors including hypermutated donors against 12 driver discovery methods in both coding and non-coding elements.

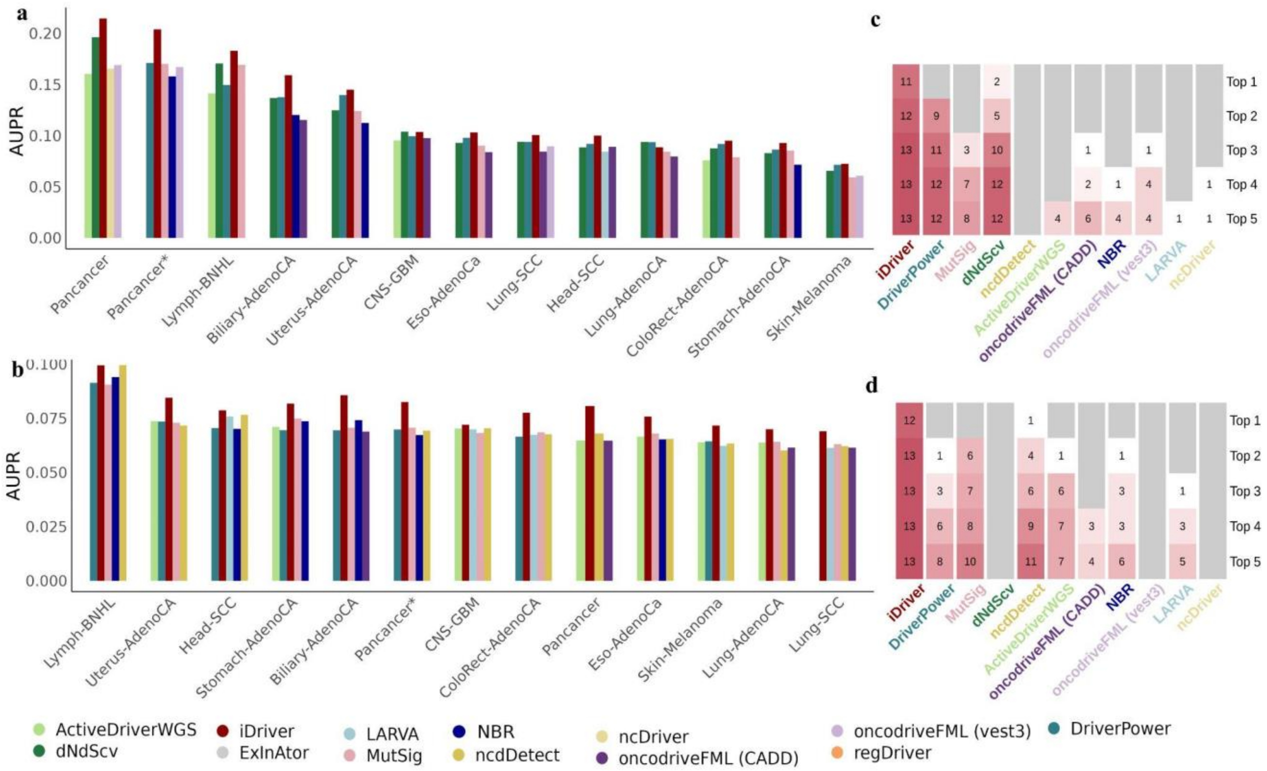


**Supplementary Figure 6** Performance comparison of iDriver on cohorts with at least one hypermutated donors, hypermutated donors included, against 12 published driver discovery methods across 13 PCAWG cancer cohorts for both coding (a and c) and non-coding (b and d) elements. The grouped bar plots (a and b) show the AUPR (Area Under the Precision-Recall curve) scores per cancer cohort for top 5 methods. Heatmaps (c and d) show how many times each method has the top-k AUPR across all cohorts (k=1, 2, ,..,5) highlighting iDriver as the top-performing method for both non-coding and coding elements in the majority of cohorts.

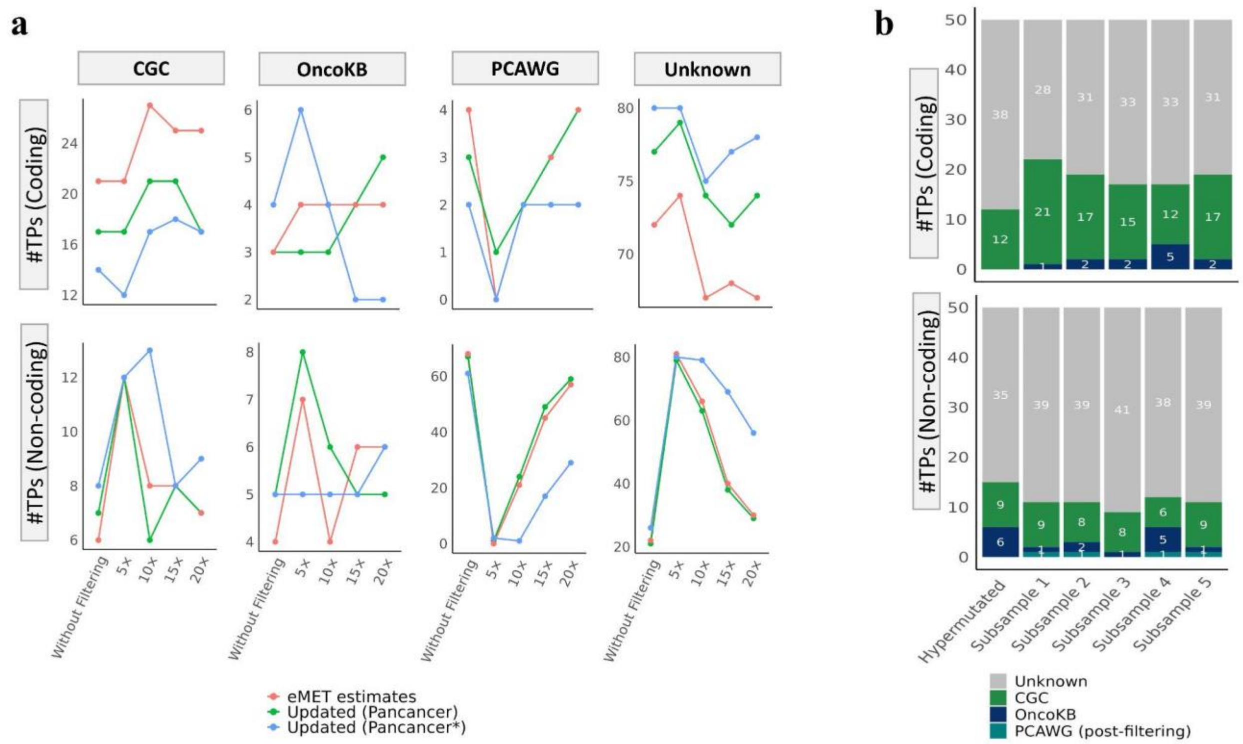


**Supplementary Figure 7** Performance of iDriver on hypermutated samples in coding and non-coding elements **(a)** Performance of iDriver under different BMR estimation strategies: i) by directly estimating mutation counts from the hypermutated cohort itself using eMET, ii) by applying eMET to the Pancancer cohort and subsequently adjusting the predicted mutation counts based on the ratio of mutation rates between the hypermutated and Pancancer cohorts, and iii) by applying eMET to the Pancancer* cohort and adjusting the predicted mutation counts using the corresponding ratio between the hypermutated and Pancancer* cohorts. The updated BMR of the Pancancer cohort yields the fewest false positives while recovering the largest number of cancer genes listed in CGC, OncoKB, or PCAWG. **(b)** Numbers of coding and non-coding elements among the top 100 iDriver hits in the hypermutated cohort and in five non-hypermutated subsamples matched to the hypermutated cohort by the number of donors per cancer type. Notably, the number of identified cancer genes in the hypermutated cohort is comparable to that in the non-hypermutated subsamples, demonstrating the capability of iDriver to identify cancer genes even when applied exclusively to a hypermutated cohort.

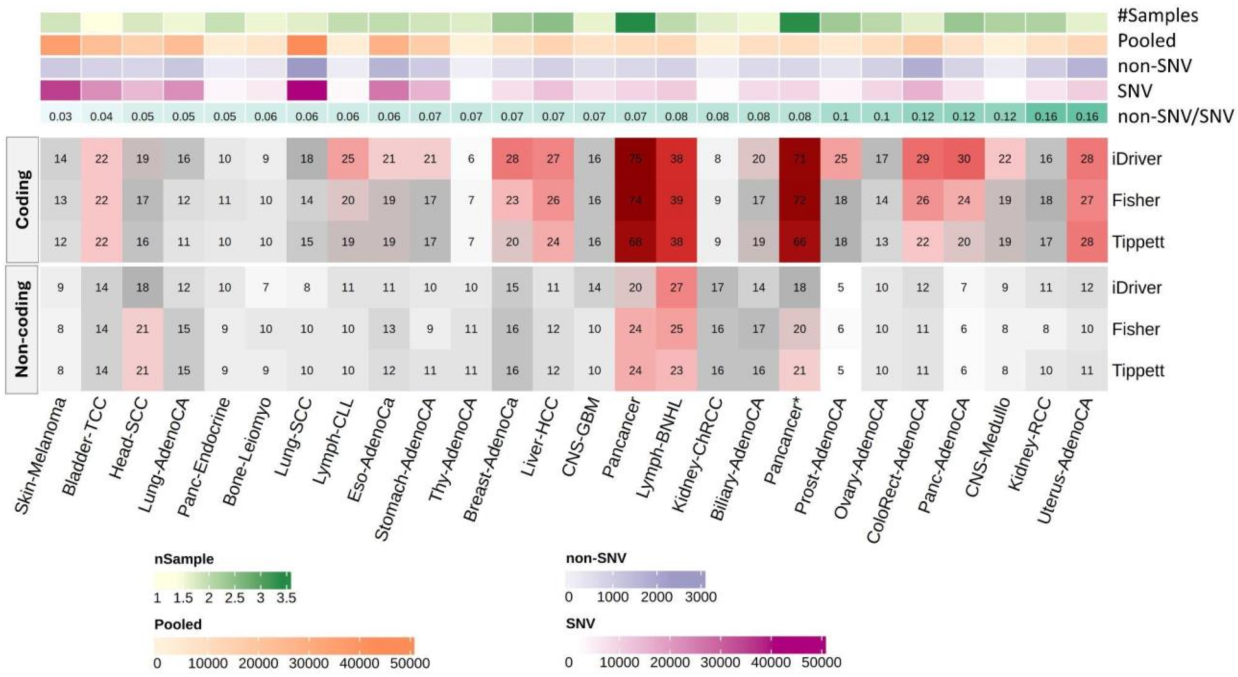


**Supplementary Figure 8** Comparison of separate modeling for SNVs and non-SNVs followed by p-value combination using Fisher's and Tippett's methods. Each cell in the heatmap shows the number of TPs in the top 100 elements for each cancer type in coding and non-coding elements using iDriver (pooled version), Tippett, and Fisher combined methods. Overall, the pooled version performs slightly better than the separate modeling approach.

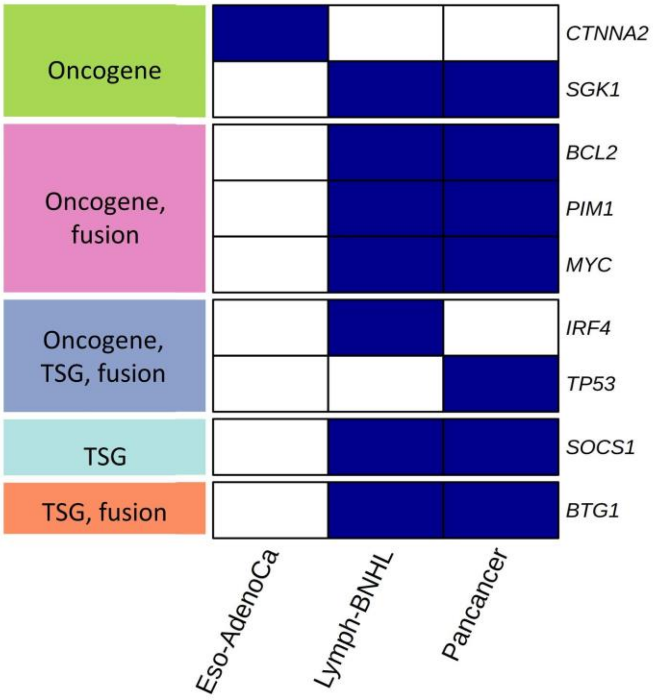


**Supplementary Figure 9** Known cancer genes among the significant synonymous hits, annotated with their roles in cancer based on CGC COSMIC.

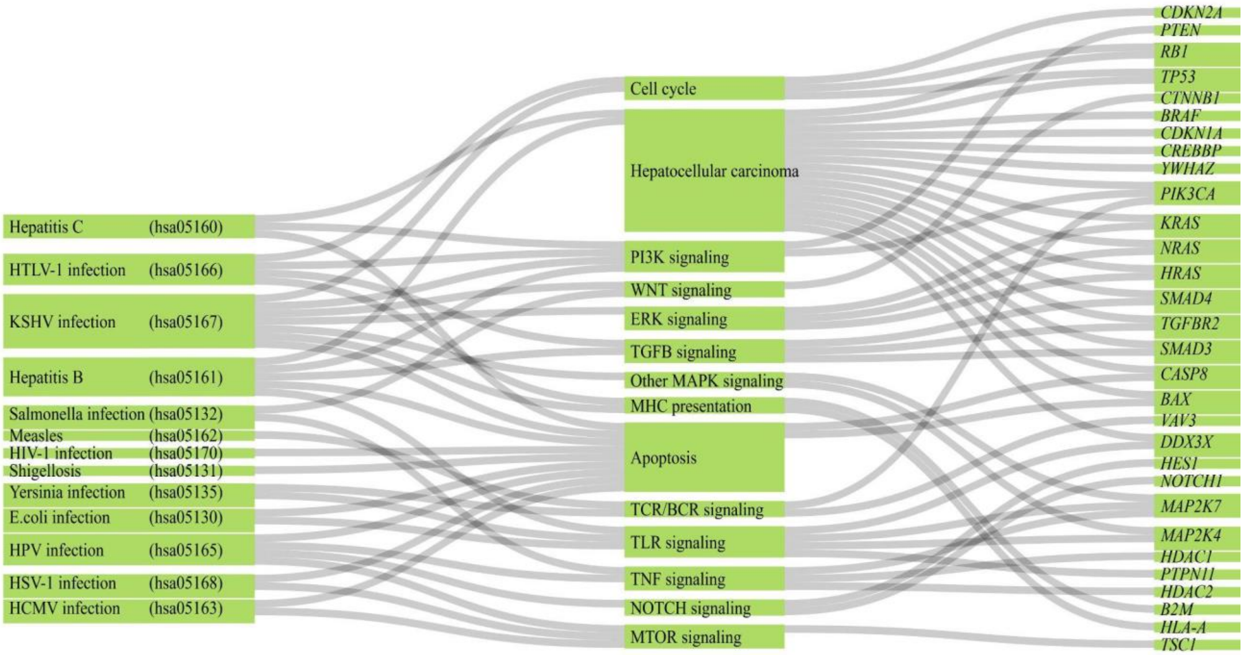


**Supplementary Figure 10** Mutated KEGG Infection related pathways and their networks associated with significant hits reidentified by iDriver. This plot shows the networks that are associated with cancer hallmarks.

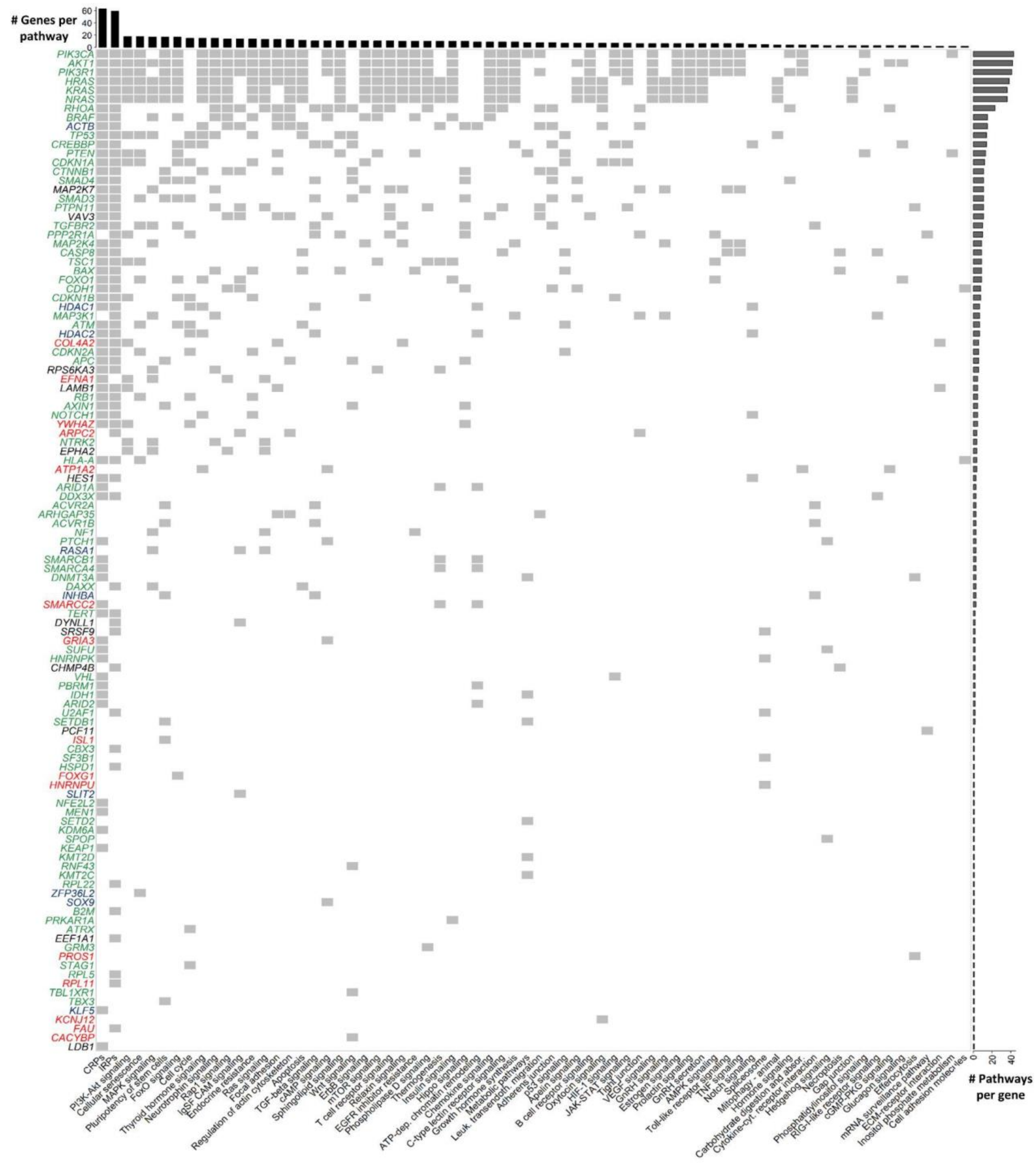


**Supplementary Figure 11** Significant hits identified by iDriver for the Pancancer* cohort and their associated pathways. Each gene may participate in multiple pathways. Gene symbol colors indicate whether the gene is listed as a COSMIC CGC gene (green), an OncoKB cancer gene (blue), a PCAWG cancer gene (black), or a novel candidate called by eMET (red).

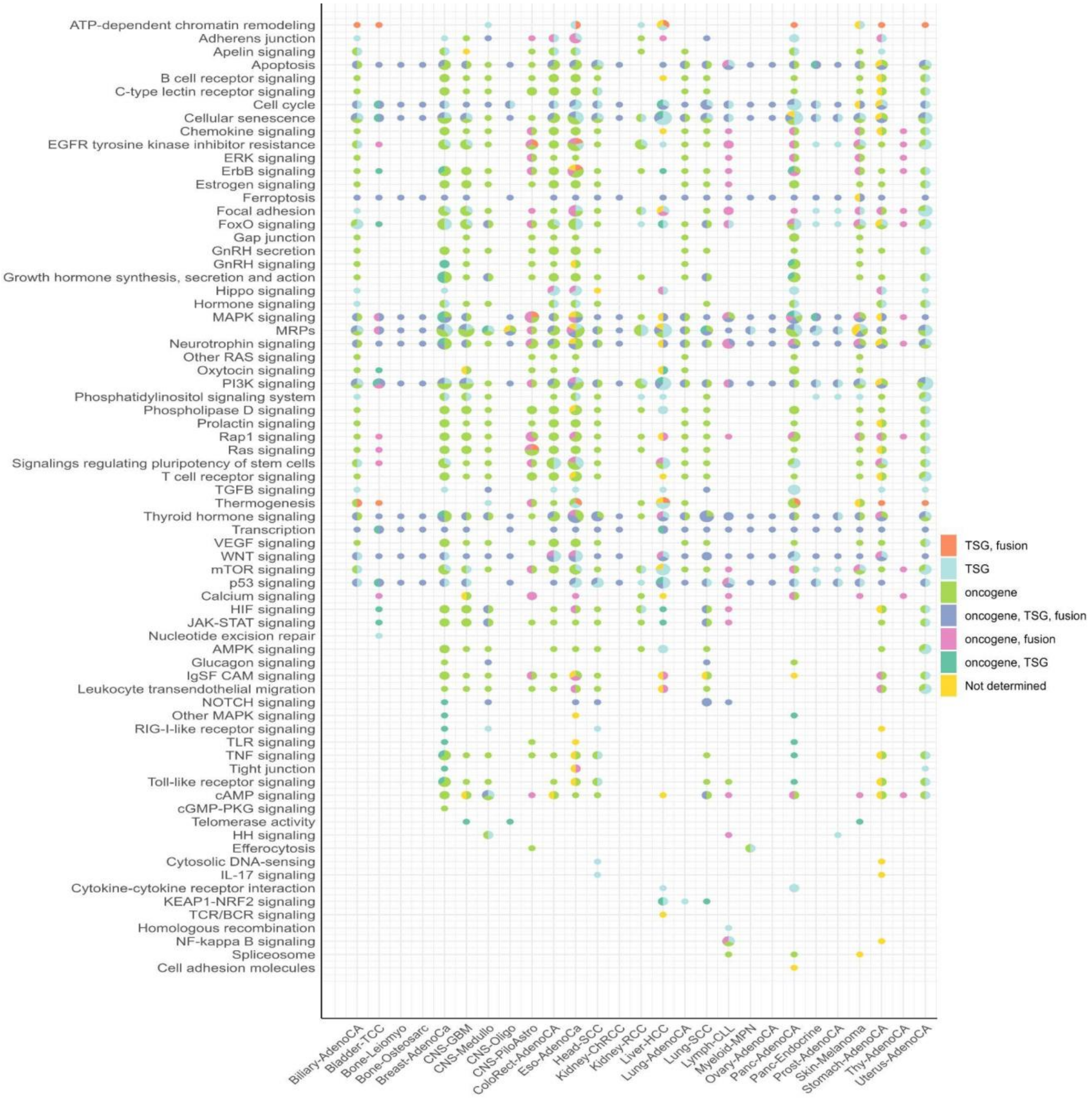


**Supplementary Figure 12** Mutated pathways across cancer types. For each cohort–pathway pair, the size of the pie chart represents the number of mutated genes, and the segments indicate the distribution of genes according to their annotated roles in cancer. This figure groups together pathways that share the same network elements across different KEGG pathways including infection related pathways.

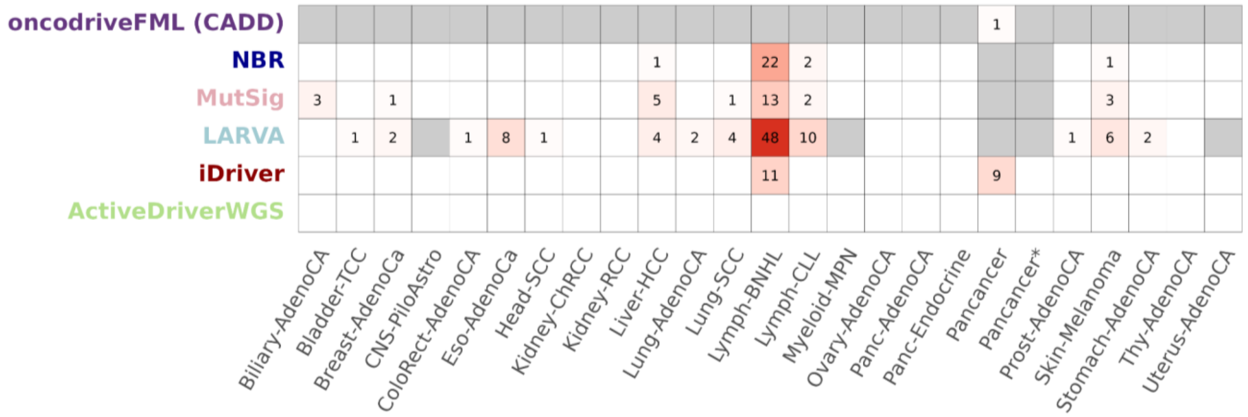


**Supplementary Figure 13** False positives reported for the simulated data across cohorts by different methods applied on all element types. White cells indicate that the method did not report any false positives for a given cohort. Grey cells indicate that either no results were available for the method in that cohort, or that the cohort had fewer than 95% of the statistical tests compared to iDriver, making the comparison unfair.

**Supplementary Table 1. Source links for the data used in this study.**

| Data | Source | Open access |
|---|---|---|
| CADD scores | https://cadd.gs.washington.edu/ | Yes |
| PCAWG_randomised_sanger.tcga.controlled.tgz | dbGaP (more information: s3://icgc25k-open/PCAWG/drivers/metadata/simulated_mutations/*) | No |
| PCAWG_randomised_sanger.icgc.controlled.tgz | ICGC DACO (more information: s3://icgc25k-open/PCAWG/drivers/metadata/simulated_mutations/*) | No |
| p-values for the type I error analyses of 12 driver discovery methods | s3://icgc25k-open/PCAWG/drivers/p-values/p-values.zip* | Yes |
| final_consensus_passonly.snv_mnv_indel.icgc.public.maf.gz | s3://icgc25k-open/PCAWG/consensus_snv_indel/final_consensus_passonly.snv_mnv_indel.icgc.public.maf.gz* | Yes |
| final_consensus_passonly.snv_mnv_indel.tcga.controlled.maf.gz | dbGap study phs000178 | No |
| callable.bed.gz | https://figshare.com/ndownloader/files/13005797 | Yes |
| PCAWG genomic elements (bed12) | https://figshare.com/ndownloader/files/12352265 | Yes |
| gene-enhancer map | s3://icgc25k-open/PCAWG/networks/map.enhancer.gene.txt.gz* | Yes |
| OncoKB™ Cancer Gene List | https://www.oncokb.org/cancer-genes | Yes |
| COSMIC Cancer Gene Census (CGC) | https://cancer.sanger.ac.uk/census | Yes |
| PCAWG drivers | https://static-content.springer.com/esm/art%3A10.1038%2Fs41586-020-1965-x/MediaObjects/41586_2020_1965_MOESM4_ESM.xlsx | Yes |

| | | |
|---|---|---|
| sample information | s3://icgc25k-open/PCAWG/donors_and_biospecimens/pcawg_sample_sheet.tsv* | Yes |

Supplementary Table 1 lists the sources of all datasets used in this study. Entries marked with an asterisk indicate data available through an Object Storage Bucket. Further details can be found at: https://docs.icgc-argo.org/docs/data-access/icgc-25k-data.
